# Human-restricted *Salmonella* exploit preassembled flagella for intracellular motility and vacuolar escape

**DOI:** 10.64898/2026.06.15.732394

**Authors:** Maximilian Lehmann, Malte Kellermann, Michael Holtmannspötter, Rainer Kurre, Michael Hensel, Marc Erhardt

**Author notes:** These authors contributed equally to this work (Maximilian Lehmann, Malte Kellermann).

## Abstract

Human-restricted typhoidal *Salmonella enterica* serovars cause systemic infections known as enteric fever, whereas most non-typhoidal serovars cause self-limiting gastroenteritis. Both groups rely on flagella-mediated motility during intestinal colonization and invasion, yet their intracellular flagellar programs appear to diverge after host cell entry. Non-typhoidal *Salmonella* are thought to rapidly silence flagellar expression intracellularly, whereas human-restricted *S*. Paratyphi A maintains flagellar gene expression and uses motility to evade xenophagy. However, the fate and function of flagella assembled before invasion have remained unclear. Here, we used fluorescence microscopy to follow preassembled flagella during epithelial cell invasion across typhoidal and non-typhoidal *Salmonella*. We show that intracellular bacteria remain flagellated during invasion across epithelial cell lines, serovars, and invasion conditions, indicating that *Salmonella* do not universally shed or lose preassembled flagella upon host cell entry. We demonstrate that internalized flagella are disassembled within *Salmonella*-containing vacuoles, whereas cytosolic flagella are targeted by autophagy. We further show that *S*. Paratyphi A uses flagella assembled before invasion to power intracellular motility and promote escape from the *Salmonella*-containing vacuole, and that intracellular motility is a phenotype shared by typhoidal *Salmonella*. Together, these findings reveal that human-restricted *Salmonella* can exploit preassembled flagella inside host cells as part of an intracellular pathogenic strategy. We propose that intracellular motility represents an adaptation to the human epithelial environment that may promote vacuolar escape and host restriction.

**Importance:** Flagella are best known for driving bacterial motility before host cell invasion, but their fate and function after intracellular entry remain poorly understood. This study shows that *Salmonella* do not universally lose their flagella during epithelial cells invasion. Instead, both typhoidal and non-typhoidal *Salmonella* serovars enter host cells with intact preassembled flagella, but only human-restricted typhoidal serovars use them for intracellular motility. This distinction separates flagellar presence from flagellar function and identifies intracellular motility as a serovar-specific behavior associated with human-restricted systemic disease. The findings suggest that typhoidal *Salmonella* have adapted to the human epithelial intracellular environment by maintaining a motile state that may promote escape from the *Salmonella*-containing vacuole and favor cytosolic lifestyle. More broadly, this work reframes flagella not only as extracellular colonization factors but also as intracellular determinants of pathogen behavior, host adaptation, and disease outcome.

## Introduction

*Salmonella* infections remain a major public health concern and impose substantial economic and societal burdens in both industrialized and low-resource settings despite continued efforts in surveillance, prevention, and treatment [1]. Based on disease presentation in humans, *Salmonella* serovars are broadly divided into non-typhoidal serovars (NTS) and typhoidal serovars (TS) [2]. NTS, including *S*. *enterica* serovars Typhimurium (STM) and Enteritidis, infect a broad range of hosts and typically cause self-limiting gastroenteritis in humans. By contrast, TS are human-adapted pathogens that cause the systemic disease enteric fever [3, 4]. The most clinically important TS are *S*. *enterica* serovar Typhi and *S*. *enterica* serovar Paratyphi A (SPA), which together account for more than 95% of reported enteric fever cases [5].

After entering the small intestine, both TS and NTS use flagella-mediated motility to reach the small-intestinal epithelium, where they adhere to and invade host cells [6]. In the intestinal lumen, *Salmonella* rapidly induces the expression of virulence factors, including genes encoded within *Salmonella* pathogenicity islands (SPIs) [7]. Flagella-driven motility supports bacterial navigation within the intestinal lumen and helps position bacteria at the epithelial surface, while flagella and other adhesins contribute to efficient attachment to epithelial cell surfaces [8, 9]. Invasion is then mediated by the SPI-1-encoded type III secretion system (T3SS) and its effector proteins, which mediate bacterial uptake into epithelial cells in a process called trigger invasion [10]. Upon host cell contact, SPI-1 effectors are translocated into the host cytosol, where they remodel the host actin cytoskeleton, induce membrane ruffling, and drive bacterial internalization into a membrane-bound compartment termed the *Salmonella*-containing vacuole (SCV) [11].

Immediately after uptake, the nascent SCV is intrinsically unstable because of the presence and activity of the SPI-1 T3SS [12, 13]. As a result, the vacuolar membrane is prone to damage, and excessive damage can lead to SCV rupture and bacterial escape into the cytosol, which ultimately leads to increased replication [14]. In NTS including STM, intracellular adaptation is accompanied by rapid shutdown of flagellar gene expression, a strategy thought to limit immune recognition [15–17]. In contrast, recent work has shown that the human-restricted serovar SPA continues to express flagellar genes after host cell entry and uses flagella-driven motility to evade recognition by the host autophagy machinery [18, 19].

However, the fate of flagella assembled before invasion remains unknown. This question is particularly relevant in human-restricted *Salmonella*, where intracellular flagella-driven motility has been linked to evasion of host autophagy [19]. Persistence or loss of preassembled flagella after entry may also shape immune detection, SCV integrity, and cytosolic adaptation in both NTS and TS. We therefore used super-resolution stimulated emission-depletion (STED) and lattice light sheet microscopy (LLSM) to determine whether *Salmonella* retain preassembled flagella during epithelial cell invasion, how these flagella are handled after internalization, and whether intracellular motility in SPA depends on flagella assembled before host-cell entry.

## Results

### Preassembled flagella are retained during *Salmonella* invasion

To enable temporal tracking of flagella by fluorescence microscopy, a rapid and reliable labeling strategy was required to differentiate old from newly synthesized flagella. Conventional antibody-based approaches involve prolonged incubation steps that are incompatible with live-cell imaging (LCI) of dynamic processes. To circumvent this, we used a maleimide-based labeling method to visualize flagella assembled before invasion [20, 21]. Maleimide conjugation generates a stable thioether bond at physiological pH (6.5–7.5) between a dye-linked maleimide group and accessible cysteine residues on the target protein [20]. Since SPA and STM FliC flagellins differ particularly within the D3 domain (SI Appendix, Fig. S 1), we introduced cysteine residues at distinct positions in D3 domains of each strain [22]. As STM strain background we used a SL1344 *fliC*-ON strain unable to express the second flagellin *fljB* by deleting the recombinase responsible for flagellin phase variation [23]. Consistent with the reported monophasic phenotype of SPA [24], our STM background strain expressed only the FliC flagellin. All engineered flagellin variants were tested for function, and no defects in swimming motility were observed relative to their respective wild-type (WT) strains (SI Appendix, Fig. S 2). HeLa WT cells were then infected with SPA FliC^E234C^ and STM FliC^E247C^ with maleimide-labeled flagella with an MOI of 30 as illustrated (Figure 1A). Using 3D-STED, we observed that intracellular STM or SPA colocalized with maleimide-labeled flagella 1 h post infection (p.i.) as demonstrated in (Figure 1B). Interestingly, detached flagellar filaments accumulated around invasion sites from SPA and STM respectively when cells were not washed after invasion (Figure 1C).

**Figure 1:**
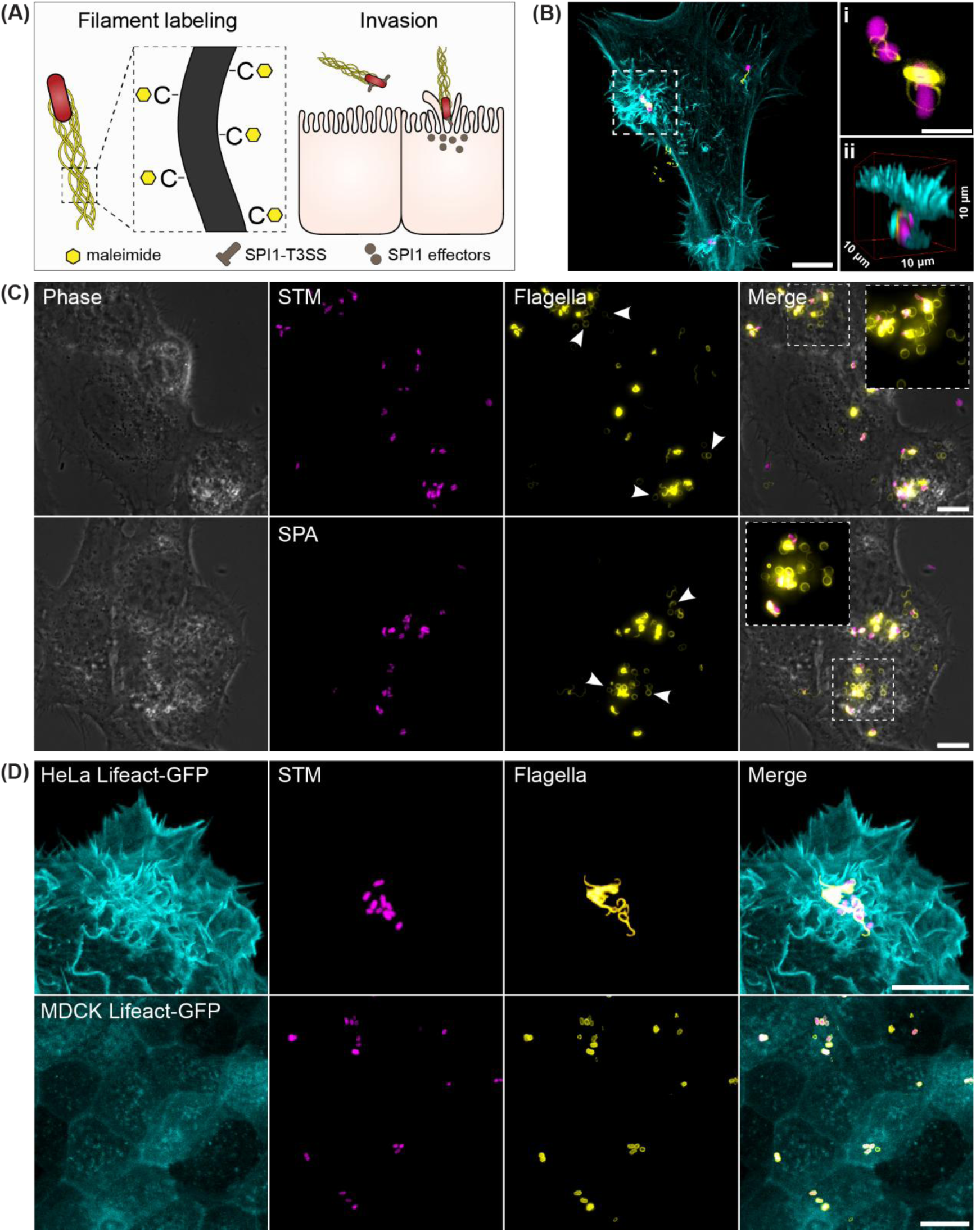
*Salmonella* retain flagella during host cell invasion. **(A)** Schematic of the flagellar-labeling approach used to monitor *Salmonella* flagellation during host-cell invasion. **(B)** HeLa LifeAct-GFP cell infected with STM FliC^E247C^, maleimide-labeled flagella and constitutive expression of mScarlet-I. Scale bar, 10 µm. Colocalization of flagella and bacteria are shown in **(i)** scale bar, 2 µm and 3D-STED of intracellular localization is demonstrated in **(ii)**. **(C)** HeLa WT cells infected with STM FliC^E247C^ and SPA FliC^E234C^ directly after invasion. Broken filaments are indicated by white arrows. Scale bars, 10 µm. **(D)** STED microscopy of HeLa LifeAct-GFP and MDCK LifeAct-GFP cells infected with STM WT constitutively expressing plasmid-encoded mScarlet-I. Flagella are visualized by immunostaining against the respective serovar-specific flagellins FliC and FljB. Scale bars, 10 µm.

To visualize the events during invasion in more detail, we used super-resolution LLSM [25]. LLSM revealed that flagellar filaments wrap tightly around the bacterial body as the bacterium is engulfed and enclosed within the nascent SCV (Movie S1 and S2). Notably, a substantial proportion of bacteria lost one or more filaments during this process, supporting the phenotype observed before (Movie S3). We infer that the extensive actin remodeling and membrane ruffling driven by SPI-1 effectors imposes mechanical forces on filaments sufficient to cause breaking of individual flagella [26, 27]. To test if intracellular flagellation is an artifact caused by flagella-maleimide labeling, and to check if flagella are also being taken up into polarized epithelial cells with brush border, we verified flagella signals by immunostaining. Therefore, HeLa and MDCK cells stably expressing LifeAct–GFP were infected with STM and SPA strains constitutively expressing mScarlet-I from plasmid and imaged using STED microscopy [28]. We found that in both HeLa and MDCK cells, intracellular STM showed flagellation at 30 min p.i. (Figure 1D), confirming that *Salmonella* serovars do not universally shed all their filaments during entry in epithelial cells with or without apical brush border [29]. Most broken filaments were no longer present, likely due to intensive washing steps. Interestingly, we observed that early after SPA invasion, the maleimide flagellar signal not only surrounded many bacteria in a pattern resembling the SCV shape, but in contrast extended beyond this vacuole-like confinement, consistent with the presence of cytosolic flagella (SI Appendix, Fig. S 3). We next examined flagellation in cytosolic STM. We used a glucose-6-phosphate (G6P)-responsive reporter to identify cytosolic STM [13]. In contrast to SPA, cytosolic STM showed no detectable flagella (SI Appendix, Fig. S 4). We then directly compared intracellular flagellation in SPA and STM. After invasion, SPA frequently displayed a higher degree of flagellation than STM (SI Appendix, Fig. S 5). This difference most likely reflects the higher level of flagellation in SPA before host-cell entry, suggesting that serovar-specific differences in flagellation before invasion contribute to the distinct intracellular phenotypes of these strains (SI Appendix, Fig. S 6). Because *Salmonella* can enter host cells through both trigger and zipper invasion mechanisms [30], we asked whether intracellular STM that entered via the SPI-1 T3SS-independent zipper mechanism also remained flagellated. To test this, we constructed a *invA* mutant to inhibit trigger invasion and introduced a plasmid constitutively expressing the *Yersinia* invasin gene *inv* to synthetically confer zipper invasion to STM (SI Appendix, Fig. S 7) [31]. We found that similar to bacteria entering by trigger invasion, intracellular STM that invaded via the zipper mechanism also retained flagella (SI Appendix, Fig. S 8)

### Internalized flagella are disassembled in acidified SCVs

Next, we examined the fate of flagella after internalization into host cells. Thus, HeLa WT cells were infected with STM WT bacteria. At 1 h, 3 h, and 6 h p.i., cells were fixed and both FliC and FljB flagellins were detected by immunostaining (SI Appendix, Fig. S 9). At 1 h p.i., flagella appeared as sharp, well-defined filaments. By 3 h p.i., the signal had weakened and the filamentous pattern had become increasingly punctate and diffuse around the bacteria. At 6 h p.i., flagellar staining was nearly absent or appeared as diffuse signal around bacteria inside SCVs. Therefore, we hypothesized that the acidic environment in vacuoles destabilizes flagella or promotes their disassembly. Consistent with this idea, we performed immunostaining against FliC of flagellated STM fliC-On after 2 h in buffers of varying pH. After wash and subsequent immunostaining, we observed a progressive reduction in flagellar signal with decreasing pH. As described before [32], after 2 h at pH 3, no flagella signal could be observed (SI Appendix, Fig. S 10). We asked if flagella disassembly could be prevented by interfering with the acidification of the SCV using the V-ATPase and autophagy inhibitor Bafilomycin A1 [33, 34]. Therefore, HeLa WT cells were infected with STM SL1344 *fliC*-On strain with anhydrotetracycline (AnTc) inducible flagella master regulator (P*_tet_*-*flhDC*) to synchronize flagellation before invasion and to prevent flagella expression after invasion. HeLa cells were either treated with a DMSO control or 100 nM Bafilomycin A1. While flagellar filaments could not be detected in the DMSO control at 8 h p.i., flagellar filaments remained detectable inside the Bafilomycin A1 treated cells, further supporting the role of acidification for flagella disassembly inside SCVs (Figure 2A). Next, we aimed to identify the fate of flagella that ended up in the host-cell cytosol, specifically for SPA infected cells. We hypothesized that flagella located in the cytosol are targeted by host autophagy for lysosomal degradation [35]. To test this, HeLa cells stably expressing the autophagic marker LC3B-GFP [36] were infected with SPA FliC^E234C^ expressing mScarlet-I and maleimide-labeled filaments. As expected, single detached flagellar filaments and bacteria-associated filaments colocalized with LC3B-GFP, demonstrating that cytosolic flagella are targeted for downstream lysosomal degradation (Figure 2B).

**Figure 2:**
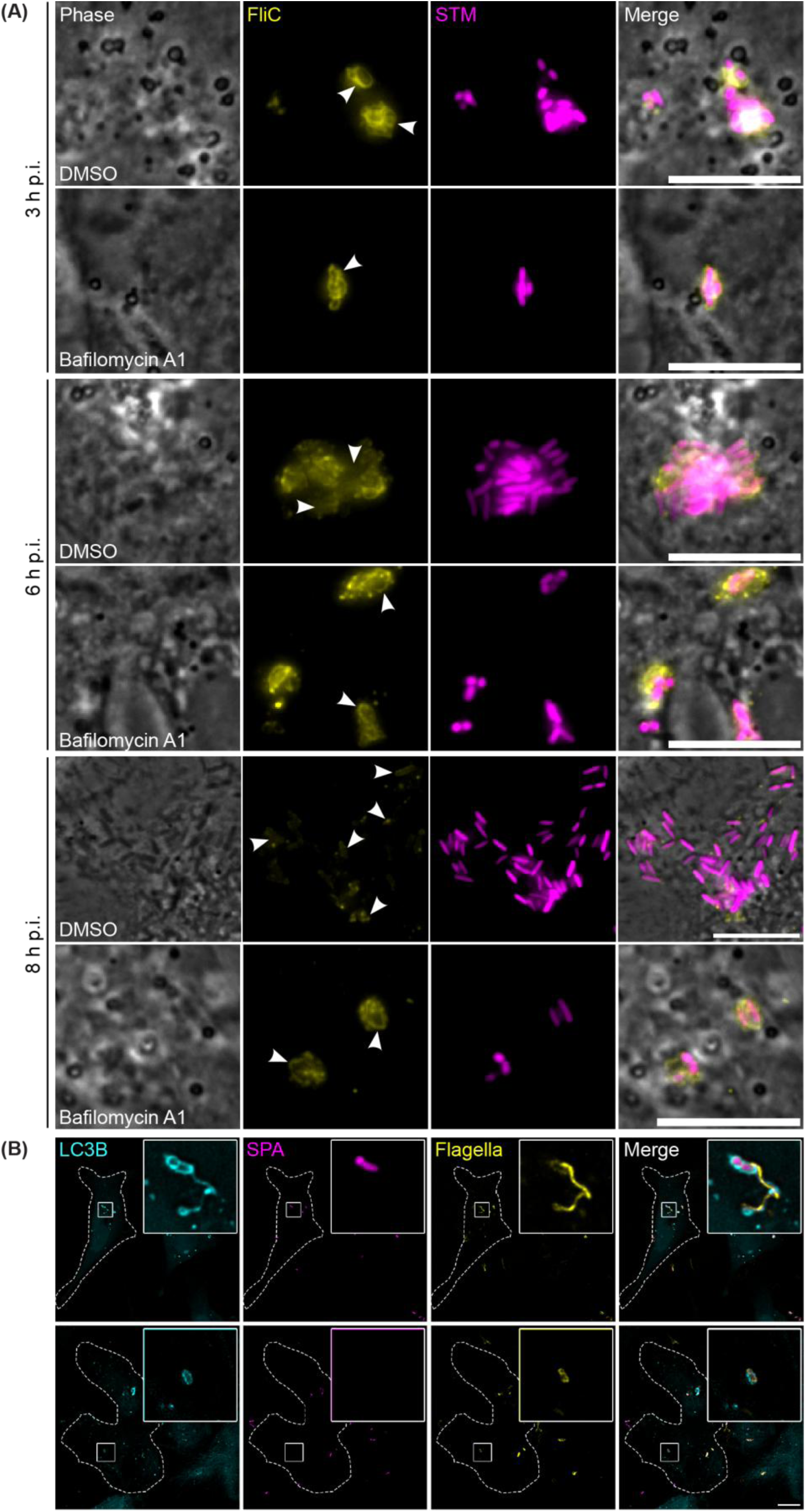
Internalized flagella degrade over time. **(A)** HeLa cells infected with STM *fliC*-On P*_tetA_*-*flhDC* expressing mScarlet-I at MOI 30 were fixed at 3, 6 and 8 h p.i. followed by immunostaining against flagellin FliC. HeLa cells were either treated with DMSO-control or 100 nM Bafilomycin A1 starting at 1 h before infection. White arrows indicate the different flagella states over time. Scale bars, 10 µm. **(B)** HeLa LC3B-GFP infected with SPA FliC^E234C^ expressing mScarlet-I and maleimide-labeled for visualization of flagella at MOI50 were imaged 1 h p.i. Scale bar: 10 µm.

### Flagellar motility promotes SCV escape by *S.* Paratyphi A

Cytosolic motility of SPA has been described previously [18]. However, during LCI of intracellular SPA and STM at early stages p.i. (1-3 h), we observed a previously undescribed form of intracellular motility. SPA displayed movement within the SCV, which we termed intra-SCV motility as movement can already be observed within LAMP1-positive compartments in HeLa cells (Figure 3A, Movie S4). This phenotype became apparent at approximately 2 h p.i., concomitant with the onset of cytosolic motility. Microscopic quantification of LAMP1-positive compartments containing either SPA or STM showed that intra-SCV motility occurred in approximately 30% of SPA-containing SCVs but was nearly absent from vacuoles harboring STM after 2 h p.i. (Figure 3B). Because intra-SCV motility occurred alongside cytosolic motility, we hypothesized that flagellar rotation within the SCV promotes escape from the vacuole. To test this idea, we generated SPA Δ*fliC* and Δ*motAB* strains in a strain background where we placed the expression of the SPI-1 master regulator *hilA* under arabinose control (SPA P*_ara_*-*hilA*), to enable synchronization of invasion (SI Appendix, Fig. S 11). Next, we introduced a dual-reporter plasmid for constitutive mScarlet-I expression together with a P*_uhpT_*-mNeonGreen cytosolic reporter with ASV-tag for accelerated degradation [37], and the *uhpABCT* operon from STM for G6P uptake. To synchronize infection, bacteria were centrifuged onto host cells for 5 min. Using flow cytometry, we quantified the fraction of infected HeLa cells containing cytosolic SPA and found that the relative proportion of cytosolic SPA on infected cells was significantly reduced in both mutants, indicating that motility contributes to SCV escape in SPA (Figure 3C.) Notably, the Δ*motAB* strain, which still produces flagella but is non-motile, yielded a higher fraction of HeLa cells containing cytosolic bacteria than the Δ*fliC* strain. Recent work has shown that the SPI-1 T3SS punctures the nascent SCV and promotes its rupture [12]. It is therefore plausible that the flagellar filament, a large surface structure of multiple µm length, composed of thousands of flagellin subunits, likewise contributes to mechanical disruption of the SCV membrane. We then sought to directly visualize motility-mediated SCV exit using LLSM. For this purpose, we used HeLa cells stably expressing HaloTag-fused to the equinatoxin domain that specifically binds sphingomyelin (EqtSM), as an infection model [38]. These reporter cells reliably indicate membrane damage resulting in sphingomyelin in cytosolic leaflet of endosomal membranes, including damaged SCVs. HeLa EqtSM cells were stained with a HaloTag ligand and infected with SPA expressing mScarlet-I and maleimide-labelled flagella. LCI captured SPA escaping from damaged SCVs after extensive intra-SCV movement (Figure 3D, Movie S5). During this process, SPA lost most of its flagella. To test whether flagella assembled before invasion contribute to intracellular motility of SPA, we infected HeLa cells with SPA, stained flagella, and imaged the cells starting at 2 h p.i., when intracellular motility becomes detectable. We found that SPA motility was mediated by flagella assembled before invasion, because motile bacteria remained associated with maleimide-labelled flagella (Figure 3E, Movie S6 and S7). Notably, motile SPA carried no more than three filaments, consistent with our previous observation of substantial flagella loss during SCV exit.

**Figure 3:**
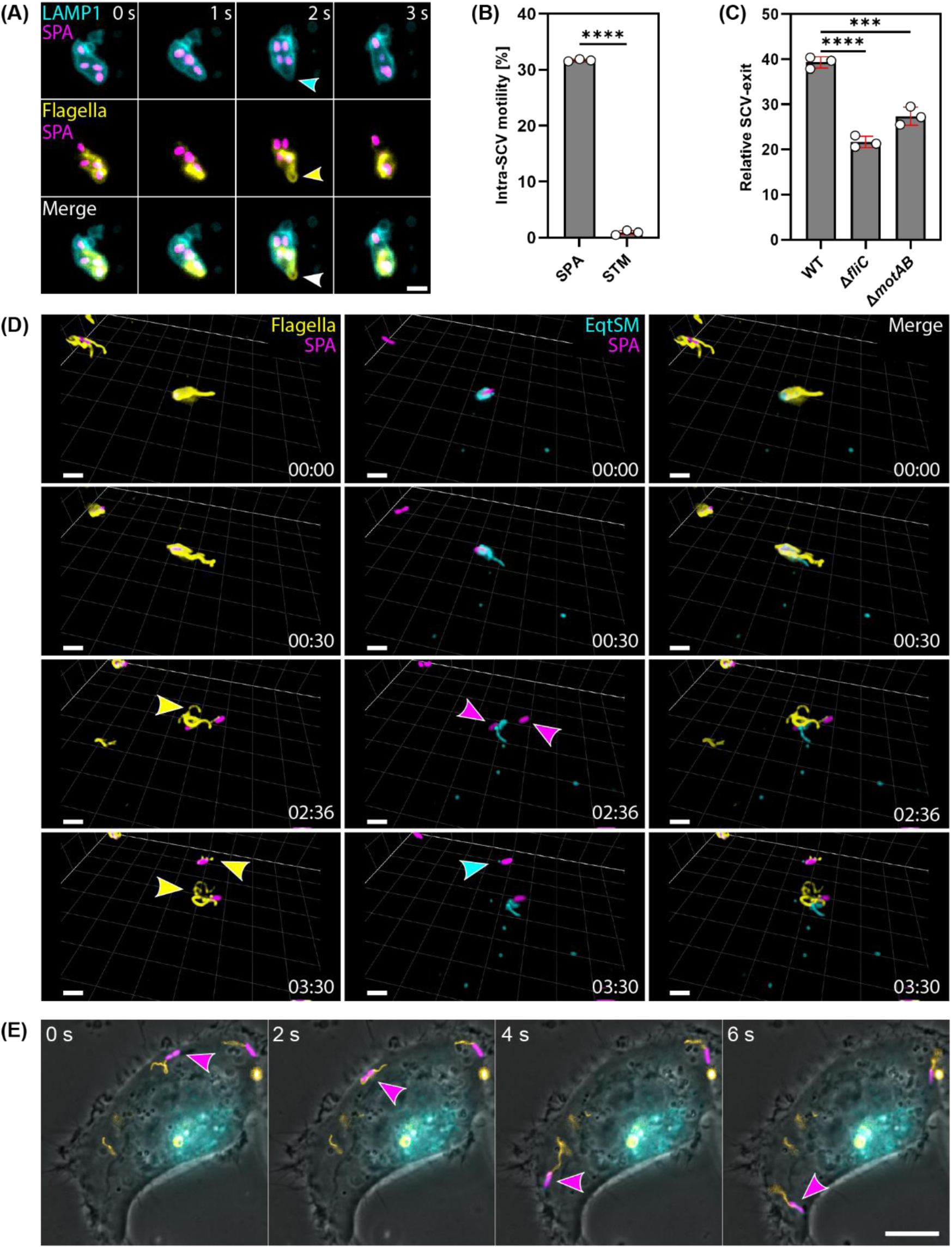
Flagella motility of SPA is linked to bacterial escape from the SCV. **(A)** HeLa cells stably expressing LAMP1-GFP were infected with SPA FliC^E234C^ expressing mScarlet-I at MOI 30 and imaged between 2 and 3 h p.i. Shown is a time-lapse panel of motile SPA with prior to invasion labeled flagella in a LAMP1-positive compartment; scale bar: 2 µm. See also Movie S4. **(B)** For quantification of intra-SCV motility HeLa LAMP1-GFP cells were infected with STM and SPA expressing mScarlet-I respectively. In total more than 600 LAMP1-positive compartments containing *Salmonella* were quantified for each serovar in three biological replicates. **(C)** Subfraction of infected HeLa cells that contain cytosolic SPA demonstrated as relative exit from total infected cells. **(D)** Flagella mediated SCV exit in HeLa EqtSM cell infected by SPA FliC^E234C^. Yellow arrows indicate breaking of flagella; magenta arrows indicate bacterial division, and cyan arrow shows exit of SPA from SCV. See also Movie S5. **(E)** A cell representative for intracellular motility of SPA by flagella assembled before invasion. Arrows indicate positions of SPA moving in cytosol. See also Movies S6 and S7.

### Intracellular motility is a shared feature of typhoidal *Salmonella*

SPA displays an intracellular motility phenotype that contrasts sharply with that of STM during host cell infection. While NTS become non-motile upon internalization, SPA retains flagellar gene expression and displays flagellum-mediated motility in the host cell cytosol. These divergent behaviors suggest that serovar-specific differences in flagellar regulation may contribute to human adaptation and systemic dissemination. To determine whether intracellular flagella-dependent motility is unique to SPA or shared by other *Salmonella* serovars, we compared the intracellular phenotypes of a panel of clinical and reference isolates. In total, 30 isolates representing 10 serovars were examined. To minimize variation arising from serovar-specific differences in invasion and to synchronize entry, all isolates were transformed with a plasmid encoding inducible expression of *hilA* under P*_tetA_* control and constitutive expression of mScarlet-I for LCI. HeLa WT cells were infected with each isolate at MOI 30, and intracellular behavior was assessed by LCI from 2 h to 3 h p.i. Strikingly, all TS tested, including isolates of SPA, *S*. Paratyphi B (SPB, systemic lineage), and *S*. Paratyphi C (SPC), exhibited both cytosolic motility and intravacuolar motility, whereas all tested NTS isolates remained non-motile under matched experimental conditions (Table 1). Notably, all tested SPB isolates of the non-systemic D-tartrate-positive variant Java [39] which causes gastroenteritis in humans, remained non-motile. These findings indicate that intracellular motility is a shared phenotype of *Salmonella* serovars capable of systemic spread in humans. Because motility has been implicated in SCV escape of SPA, we hypothesized that TS would exit the vacuole more frequently than NTS. To test this, we quantified cytosolic subfraction in HeLa cells for three NTS and three TS isolates (SI Appendix, Fig. S 12). Flow cytometry showed clearly that SPA has the highest apparent frequency of cytosolic exposure among the isolates tested. However, no consistent trend distinguishing TS from NTS was observed.

**Table 1:**
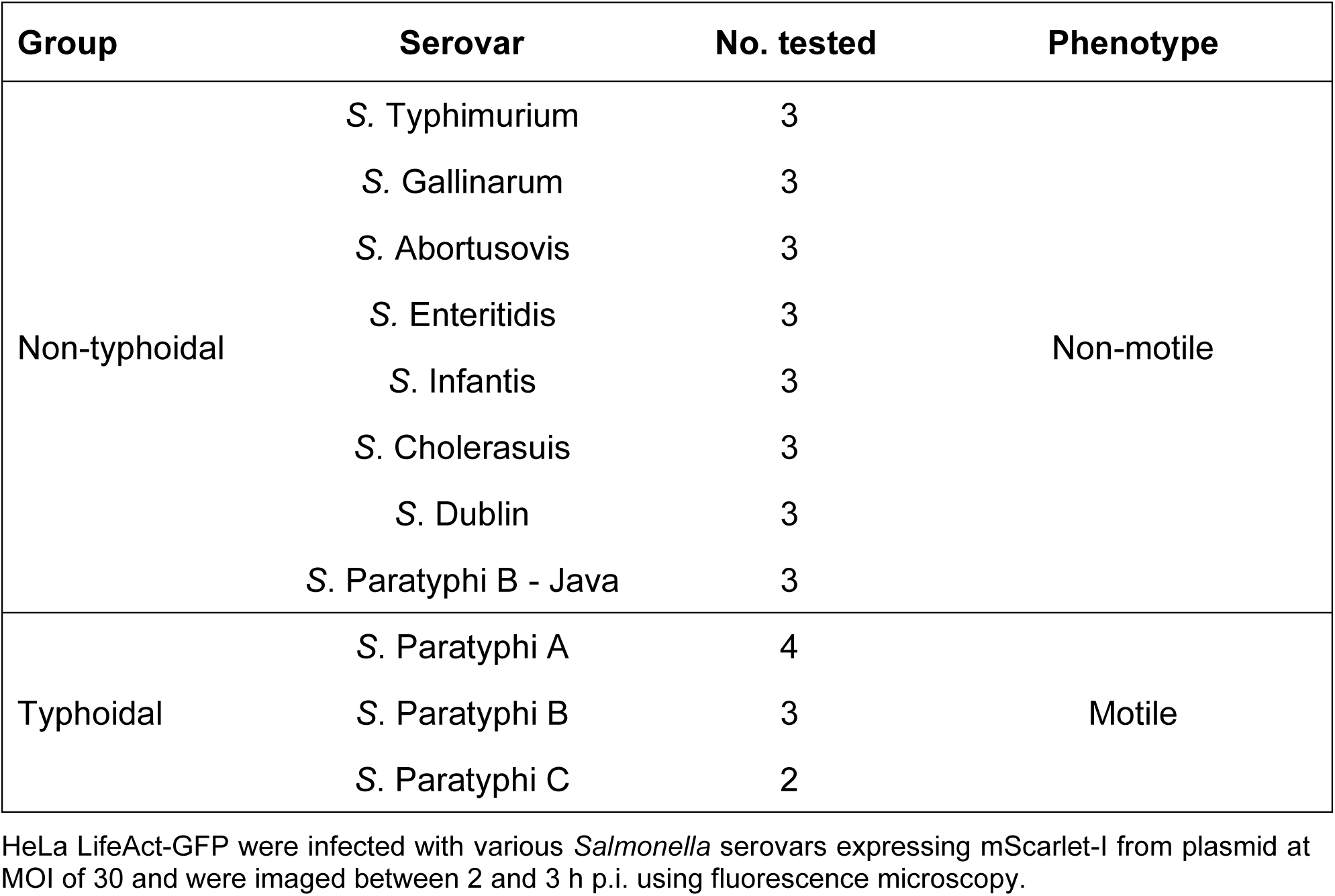
Intracellular motility is a shared feature of typhoidal *Salmonella*.

| Group | Serovar | No. tested | Phenotype |
| --- | --- | --- | --- |
| Non-typhoidal | *S. Typhimurium* | 3 | Non-motile |
|  | *S. Gallinarum* | 3 |  |
|  | *S. Abortusovis* | 3 |  |
|  | *S. Enteritidis* | 3 |  |
|  | *S. Infantis* | 3 |  |
|  | *S. Cholerasuis* | 3 |  |
|  | *S. Dublin* | 3 |  |
|  | *S. Paratyphi B - Java* | 3 |  |
| Typhoidal | *S. Paratyphi A* | 4 | Motile |
|  | *S. Paratyphi B* | 3 |  |
|  | *S. Paratyphi C* | 2 |  |

HeLa LifeAct-GFP were infected with various *Salmonella* serovars expressing mScarlet-I from plasmid at MOI of 30 and were imaged between 2 and 3 h p.i. using fluorescence microscopy.

We therefore assessed the responsiveness of the cytosolic exposure reporter in each tested isolate to G6P. Reporter induction varied substantially across isolates, indicating that the P*_uhpT_*-mNeonGreen-ASV reporter is highly informative within a given strain background but less suitable for direct quantitative comparisons of SCV escape across diverse *Salmonella* serovars (SI Appendix, Fig. S 13). In addition, endpoint measurements such as flow cytometry may be influenced by isolate-specific differences in bacterial replication after host cell exit, which can affect the measured fluorescent population independently of the initial escape event. Together, these findings define an important limitation of fluorescence-based endpoint comparisons across heterogeneous clinical isolates. Complementary approaches with strain-normalized reporter calibration or time-resolved single-cell measurements will be required to resolve serovar-specific differences in cytosolic access with greater precision.

## Discussion

NTS including STM are known to downregulate flagellar gene expression soon after host cell invasion, a response thought to limit immune recognition [15–17, 40]. In contrast, the human-restricted serovar SPA maintains flagellar gene expression and displays cytosolic motility in epithelial cells, which has been linked to evasion of xenophagy [18, 19, 41]. However, the fate of flagella present before invasion had remained unresolved. By tracking flagella during invasion in SPA and STM, we found that intracellular *Salmonella* remained flagellated irrespective of epithelial cell line, serovar, or invasion mode tested. These findings argue against rapid flagellar shedding or universal loss, as described for other bacterial species [42, 43], and indicate that the disappearance of motility in STM cannot be explained by immediate loss of preassembled flagella.

Our results further show that intracellular flagella-mediated motility is not restricted to SPA but is shared among the TS tested, whereas all examined NTS isolates remained non-motile. This pattern points to a serovar-specific intracellular behavior associated with systemic spread in humans. One possible explanation is that the selective pressure to silence flagella after invasion differs in human epithelial cells. In line with this, previous work has shown that flagellin specific NAIP/NLRC4 inflammasome signaling is impaired in human intestinal epithelial cells [44, 45] and that epithelial immune responses rely primarily on LPS-mediated inflammasome rather than the recognition of flagellins or SPI-1 T3SS [46]. Together, these findings are consistent with the idea that intracellular flagella expression in human epithelial cells imposes a lower inflammasome-related cost than in other hosts, including mice [47]. In this context, maintenance of intracellular motility from human-restricted serovars may favor a cytosolic lifestyle in human epithelial cells only.

Thus, flagella expression and motility most likely also contributes restriction to hosts that lack NAIP/NLRC4 inflammasome signaling in epithelial cells [41] and could also further indicate why there are only so few TS described [2].

This interpretation is further supported by the comparison between the systemic SPB and the closely related D-tartrate-positive variant Java [39, 48]. Despite their close relationship, only the human-restricted SPB isolates displayed intracellular motility, whereas the gastroenteritis-associated Java isolates remained non-motile. Likewise, STM was non-motile shortly after invasion, whereas SPA exhibited motility within a substantial fraction of SCVs. Because motility promotes SCV escape and increases the cytosolic bacterial subpopulation in SPA, these observations support a model in which human-restricted *Salmonella* are adapted to exploit intracellular motility as part of their pathogenic strategy rather than remaining primarily vacuolar. Currently, our comparison of SCV escape across serovars is limited by the heterogeneous performance of the P*_uhpT_*-mNeonGreen-ASV reporter, which was not sufficiently robust for quantitative comparisons between isolates. Additional approaches will therefore be needed to define more precisely how intracellular motility, vacuolar escape, and cytosolic adaptation are linked across serovars.

Overall, our findings indicate that generally *Salmonella* invade host cells with their flagellar filaments attached and become internalized in SCVs, whereas the use of flagella for intracellular motility is a characteristic of human-restricted serovars. We propose that this motile intracellular behavior reflects adaptation to the environment of human epithelial cells and may contribute to host restriction and systemic dissemination.

## Methods

### Strains, Media and Growth conditions

Bacterial strains and genotypes are listed in (SI Appendix, Table S 1). *Salmonella* serovars were grown in LB (Miller) (10 g/L tryptone, 5 g/L yeast extract, 10 g/L NaCl) at 37 °C in a roller drum unless stated otherwise. Overnight (o/n) cultures were prepared by inoculating single colonies in LB medium. Day cultures were prepared by diluting o/n cultures 1:100 and cultures were grown to mid-exponential phase, if not stated otherwise. Antibiotics were added if needed for plasmid maintenance. For induction of gene expression 10 ng/ml AnTc, 0.05% - 0.4% arabinose or 0.2% G6P were added.

### Bacterial strain construction

Bacterial mutant strains were generated by λ Red recombineering for insertion of the KanSceI resistance cassette amplified from template DNA using oligonucleotides listed in (SI Appendix, Table S 1). Insertion or replacement of KanSceI cassette with respective desired genotype was verified by colony-PCR and sequencing.

### Swimming motility assay

To assess the motility phenotype of *Salmonella* strains containing FliC cysteine mutations, soft-agar (0.3% w/v bacto agar) plates were inoculated with 2 µl of an o/n culture of the tested strains. The inoculated plates were incubated at 37 °C for 4.5 h. Subsequently, the plates were scanned using the Perfection V800 Photo scanner (Epson). The diameter of the motility halo was measured using ImageJ and used as a representative of the swimming behavior of the cells.

### Host cell culture and infection

Cell lines and genotypes are listed in (SI Appendix, Table S 1). HeLa cell lines were cultured in high-glucose Dulbecco’s modified Eagle’s medium (DMEM) supplemented with GlutaMAX (Gibco) and supplemented with 10% inactivated fetal calf serum (iFCS, Sigma). MDCK cells were cultivated in MEM (+ Earle’s Salts + 4 mM Glutamax) supplemented with 10% iFCS and 1% NEAA. All cells were cultured at 37 °C in an atmosphere containing 5% CO_2_ and absolute humidity. For cell passaging, cells were washed thrice in pre-warmed (37 °C) PBS and incubated with accutase for 5 to 30 min until all cells were detached from the culture flask. To stop the detachment reaction, the same volume of DMEM (10% FCS) was added. An appropriate number of cells were passaged into a fresh culture flask. For infection experiments, bacteria were either added directly to the cells or the medium was replaced with prior prepared infection mixes followed by a 30 min incubation at 37 °C and 5% CO_2_, if not stated otherwise. After incubation, cells were treated in Gentamicin protection assays as described before. Between all steps, cells were washed thrice with pre-warmed PBS, if not stated otherwise.

### Infection experiments and microscopy

HeLa WT or stably expressing LifeAct-GFP, LAMP1-GFP or LC3B-GFP cells were seeded into ibidi treat 8-well dishes or on glass cover slips 24 h before infection. Cells were grown to 80% confluency on the day of infection. MDCK LifeAct-GFP cells were seeded 7 days before invasion to reach 100% confluence.

To determine the flagella fate, cells were infected with either SPA FliC^E234C^ or STM FliC^E247C^ cultured under 3.5 h aerobic (STM) or 16 h microaerobic (SPA) conditions. Bacterial cultures were diluted to OD of 0.2 in 1 ml PBS. Maleimide StarRed (Aberrior) was added to get a final concentration of 10 µM from 10 mM stock (DMF). Flagella were stained for 10 min at 37 °C at 300 rpm shaking and washed once before adding bacterial suspensions directly to the cells. After gentamycin protection assay, samples were than imaged with a Nikon Eclipse Ti2 inverted microscope equipped with CFI Plan Apochromat DM 60x Lambda oil Ph3/1.40 objective (Nikon) and an Orca-Fusion BT digital camera C15440, exposure 100 ms phase contrast, 100 ms 5% mScarlet-I and 100 ms 40% StarRed.

To quantify the intra-SCV motility phenotype, HeLa LAMP1-GFP cells infected with SPA FliC^E234C^ or STM FliC^E247C^ with maleimide labelled flagella were imaged at 2 h p.i. with a Nikon Eclipse Ti2 inverted microscope using triggered acquisition mode at 37°C. Triggered acquisition settings include 10 ms 10% GFP laser and 10 ms 5% mScarlet-I laser for 10 sec time lapse movies. More than 200 LAMP1 compartments surrounding bacteria were quantified for movement for each biological sample. Statistical analysis was performed in GraphPad Prism.

To test if *Salmonella* isolates share intracellular motility phenotype, isolates harboring pEM19802 (pKH70 P*_tet_*-*hilA* P*_rpsM_*-mScarlet-I, CmR) were grown for 3.5 h under aerobic conditions in LB supplemented with 10 ng/ml AnTc. Bacteria were added at MOI 30 and centrifuged onto HeLa LifeAct-GFP cells at 500 x g for 5 min and were further incubated for 25 min at 37 °C with 5% CO_2_. Samples were imaged between 2 and 3 h p.i. using triggered acquisition at 37°C with the following settings: 10 ms 10% GFP laser and 10 ms 5% mScarlet-I laser for 10 sec time-lapse movies. Movies were analysed in ImageJ.

For intracellular flagella immunostaining samples were fixed with 4% formaldehyde for 15 min and washed three times with PBS and incubated in blocking solution (2% goat serum, 2% BSA and 0.1% saponin in PBS) for 30 min. Blocking solution for polarized MDCK LifeAct-GFP cells was supplemented with 0.5% Triton X-100. Next, samples were stained with primary antibodies overnight at 4 °C (1:500) Anti-FliC STM (Difco *Salmonella* H Antiserum i) and (1:500) Anti-FljB STM (Difco *Salmonella* H Antiserum Single Factor 2) or (1:500) SPA (Anti-*Salmonella* H:a; Sifin TR1401). Accordingly, secondary antibodies were selected and samples were incubated for 1 h (1:10,000) at RT. Samples were imaged with Abberior STED Facility Line equipped with a UPLSAPO 100x oil objective, NA 1.4, a confocal Pulsed Diode Laser PDL-T 488 (10% power, line accumulation 3) for imaging the LifeAct-GFP and a Pulsed Diode Laser PDL-T 640 (power 12%, line accumulation 3) and STED laser-40-3000-775-B1R (power 12%, line accumulation 3) for imaging maleimide STAR RED. The pinhole size was set to 0.8–1.0 AU, the pixel dwell time to 5 µs and pixel size to 40 × 40 × 80 nm (xyz). Intervals for Z-stack images were set to 50 nm. Images were further processed with ImageJ.

Lattice Light-Sheet Microscopy (LLSM) was performed using a custom build system based on the original design by the Eric Betzig group [25]. Stably transfected Hela cells were seeded on plasma-cleaned and PLL-PEG-RGD-coated 5 mm round glass coverslips (Art. No. 11888372, ThermoFisher Scientific) and grown in DMEM medium. Cells expressing Equinatoxin-HaloTag (EqtSM-HaloTag) were incubated with 100 nM HTL-JFX554 for 30 min at 37 °C and infected with SPA FliC^E234C^ pEM8731 (pKH70 P*_rpsM_*-mNeonGreen, Amp^R^) that was subcultured and grown under microaerobic conditions for 16 h. To enhance bacterial contact to the host cell, centrifugation for 5 min at 500 x g was performed and followed by 25 min incubation at 37 °C with 5 % CO_2_.

After that, cells were washed three times with PBS and inserted into a custom-built sample holder, which was mounted onto a piezo stage (sample piezo) for fast sample scan imaging. This ensured that the coverslips were positioned correctly between the excitation and detection objectives inside the sample bath. The latter contained 6 ml of DMEM at a temperature of 37 °C. Humidity and temperature were controlled by a fully automatic incubation system during the entire experiment (H301-LLSM-SS316 & CO2-O2 Unit BL, Okolab, Italy). For illumination, a dithered square lattice pattern generated by multiple Bessel beams using an inner and outer numerical aperture of the excitation objective of 0.48 and 0.55, respectively, was used. As illumination sources, a 488 nm laser (2RU-VFL-P-300-488-B1R; MPB Communications Inc., Pointe-Claire, Canada) was used for SPA FliC^E234C^ pEM8731 or lifeact-SG, a 561 nm laser (2RU-VFL-P-2000-561-B1R; MPB Communications Inc., Pointe-Claire, Canada) was used for SL1344 (P*_rpsM_*-mScarlet-I) or EqtSM-HaloTag-JFX554 and a 642 nm laser (2RU-VFL-P-2000-642-B1R; MPB Communications Inc., Pointe-Claire, Canada) was used for abberior STAR RED. The final lattice light-sheet was generated by a water-dipping excitation objective (54-10-7@488-910 nm, NA 0.66, Special Optics), while emitted photons were collected by a water-dipping detection objective (CFI Apo LWD 25XW, NA 1.1, Nikon). Emission was then split via a beamsplitter (FF640-FDi01-25x36, Semrock), filtered by a corresponding dual bandpass filter (Brightline HC 523/610, Semrock) for Lifeact-SG, SPA FliC^E234C^ pEM8731, EqtSM-HaloTag-JFX554 and SL1344 (P*_rpsM_*-mScarlet-I) as well as a long-pass filter (647 LP Edge Basic, Semrock) for abberior STAR RED labelled flagella. Split emission was detected on individual sCMOS cameras (ORCA-Fusion, Hamamatsu, Japan) with a final pixel size of 103.5 nm. For image acquisition, a sequential three-channel image stack was acquired in sample scan mode by scanning the sample through a fixed light sheet with a step size of 450 nm, which is equivalent to ∼244 nm slicing with respect to the z-axis considering the sample scan angle of 32.8°. Raw data were further processed by usage of an open-source LLSM post-processing utility called LLSpy (https://github.com/tlambert03/LLSpy), which was used for deskewing, deconvolution, channel registration, and transformation. Deconvolution was performed using experimentally measured point spread functions (PSFs), obtained from 100 nm FluoSpheres™ (Art. No. F8801 and F8803, Thermo Fisher Scientific) for 488 nm and 561 nm excitation, while 170 nm PS-Speck™ (Art. No. P7220, Thermo Fisher Scientific) were used for 642 nm excitation. Deconvolution was carried out using the Richardson–Lucy algorithm with 10 iterations. Prior to acquisition, point spread functions were optimized using F8803 FluoSpheres™ and a custom-built adaptive optics system based on the AO KIT BIO from Imagine Optic. For channel alignment, registration and transformation, 200 nm sized fluorescent TetraSpeck™ microspheres (Art. No. T7280, ThermoFisher Scientific) were imaged with a step size of 350 nm. Channel transformation was performed using a two-step approach, applying an affine transformation in the xy-plane and a rigid transformation along the z-axis. Three-dimensional rendering was performed using Imaris 9.5 (Bitplane).

### Infection for full spectrum flow cytometry

HeLa cells were seeded in 24-well plates 24 h before infection to reach 100% confluence. *Salmonella* strains carrying the indicated reporter plasmids were grown o/n in LB-Miller supplemented with ampicillin or chloramphenicol at 37 °C under aerobic or microaerobic conditions. The following day, bacterial cultures were diluted 1:31 in fresh LB-Miller supplemented with ampicillin or chloramphenicol and grown 3.5 h at 37 °C. Bacteria were diluted to an OD_600_ of 0.2 and infection mixes containing bacteria were prepared and added to respective wells. Infection was performed as indicated above. Flow cytometry was performed on a Cytek Aurora full-spectrum flow cytometer using Cytek Aurora CS 5L System. Single-stained controls, unstained cells, uninfected cells, bacteria-only samples, and fluorescence-minus-one controls were included where appropriate. Instrument performance was verified before acquisition according to the manufacturer’s recommendations. Spectral unmixing was performed using single-color controls acquired with the same instrument settings as the experimental samples. Events were acquired using Spectroflow, and at least 10,000 host-cell events were recorded per sample if not stated otherwise.

Data were analyzed using FlowJo or python based FlowKit. Host cells were identified based on FSC-A and SSC-A, followed by exclusion of debris and doublets using FSC-H/FSC-A and SSC-H/SSC-A gating. Infected cells were identified based on bacterial mScarlet-I fluorescence, and reporter activity was quantified from the mNeonGreen reporter signal within the infected-cell population. Where applicable, bacterial debris or extracellular bacteria were excluded. The percentage of infected cells, median fluorescence intensity, and/or mean fluorescence intensity were calculated for each sample. Data are shown as mean from independent biological replicates.

### Flagella pH sensitivity assay

To test flagella stability at different pH levels, STM fliC-On was grown o/n, diluted 1:100 in fresh LB, and grown for an additional 2 h. The bacterial culture was then adjusted to an OD_600_ of 0.2 in 1 mL of the respective buffer and incubated for a further 2 h at 37 °C with shaking. The following buffers were used: 50 mM MOPS buffer, pH 7.0; 50 mM acetate buffer, pH 5.0; 50 mM acetate buffer, pH 4.0; and 50 mM citrate buffer, pH 3.0. After buffer incubation, bacteria were centrifuged at 2,500 × g for 10 min and washed in PBS (pH 7). Bacterial suspensions were loaded into poly-L-lysine-coated flow cells, inverted, and incubated for 2 min. Cells were then fixed with 4% formaldehyde for 5 min, followed by immunostaining against FliC. Samples were blocked in 10% BSA for 30 min. Primary antibody Anti-FliC STM (Difco *Salmonella* H Antiserum i) was incubated o/n at 4 °C followed by respective secondary antibody 1:10,000 for 1 h at RT. Samples were imaged using a Nikon Eclipse Ti2 inverted microscope.

## Acknowledgments

M.E. and M.He. acknowledge funding from the Deutsche Forschungsgemeinschaft (DFG; German Research Foundation), research grant no. 446414114, within the framework of the DFG Priority Programme SPP 2225 “EXIT Strategies of Intracellular Pathogens,”. M.E. acknowledges supported from the Max Planck Society through a Max Planck Fellowship. M.H. further acknowledges support supported by the DFG by the SPP 2225 imaging platform (research grant no. 446463289), and through SFB 1557, project P8 (research grant no. 467522186). We thank members of the Erhardt and Hensel labs for helpful discussions and critical comments on the manuscript, and Heidi Landmesser, Raúl Trepel-Washington, Monika Nietschke und Ursula Krehe for technical assistance.

## Supplementary Material

**SI Appendix, Fig. S 1:**
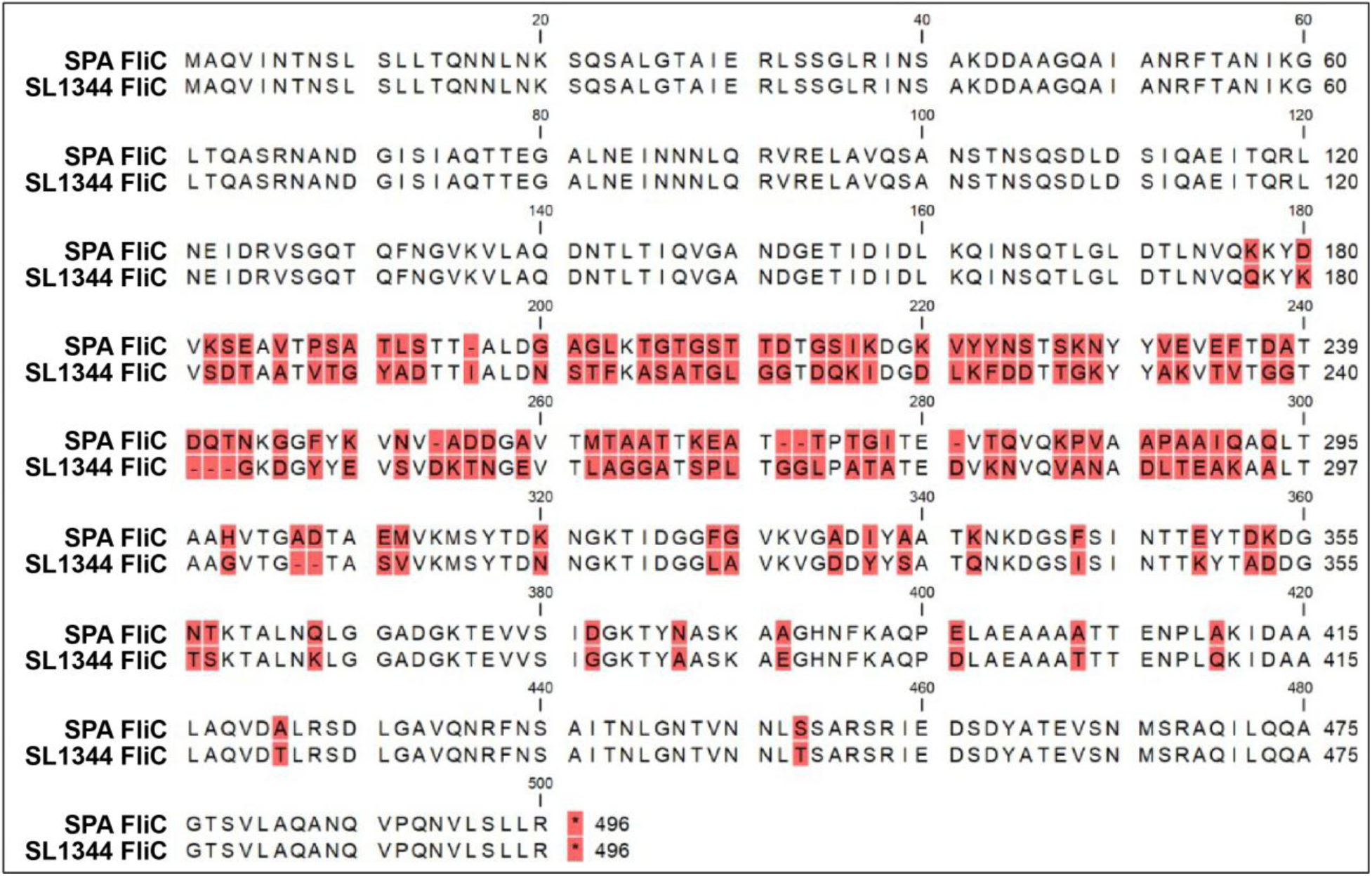
AA sequence alignment of FliC from SPA and STM SL1344. The flagellin FliC is 78.57% conserved. Differences are marked in red. AA alignment was performed in CLC Main Workbench 23.

**SI Appendix, Fig. S 2:**
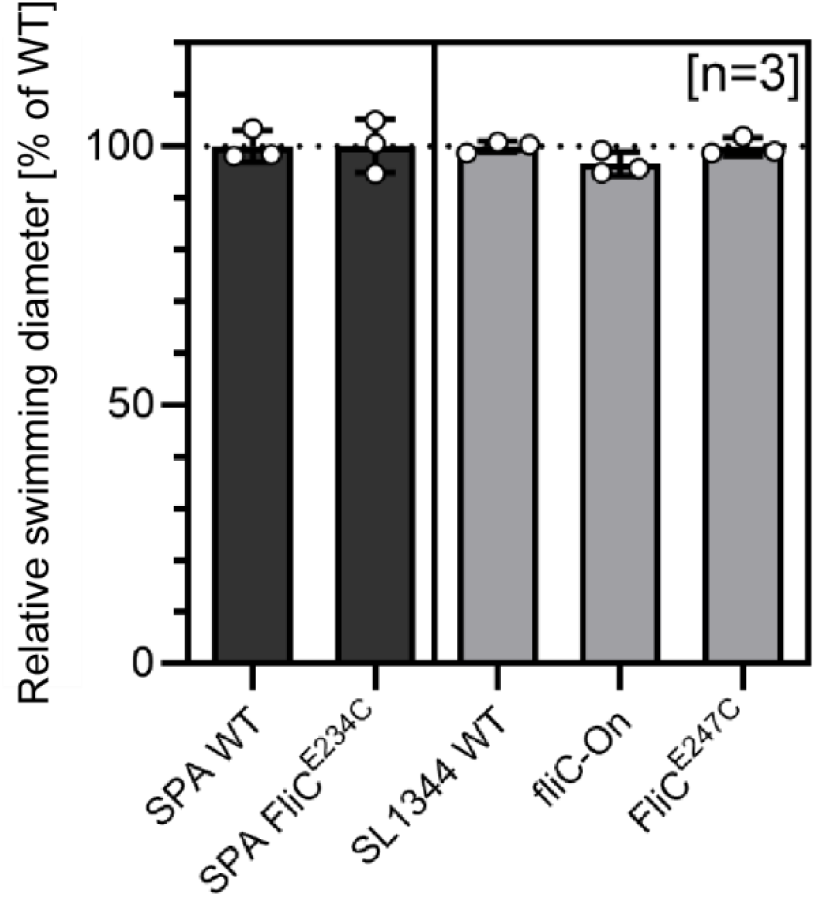
FliC cysteine exchanges do not interfere with motility. Bacteria were cultured o/n and subsequently 2 µl were inoculated into motility agar plates. Three biological replicates consisting of three technical replicates were measured and analysed after 4 h incubation at 37 °C. The diameter of the swimming halos formed by the bacteria was measured and results are presented as mean values of each biological replicate. The graph illustrates the relative motility of mutant strains compared to SPA WT or SL1344 WT, with the respective WT strain set as 100%.

**SI Appendix, Fig. S 3:**
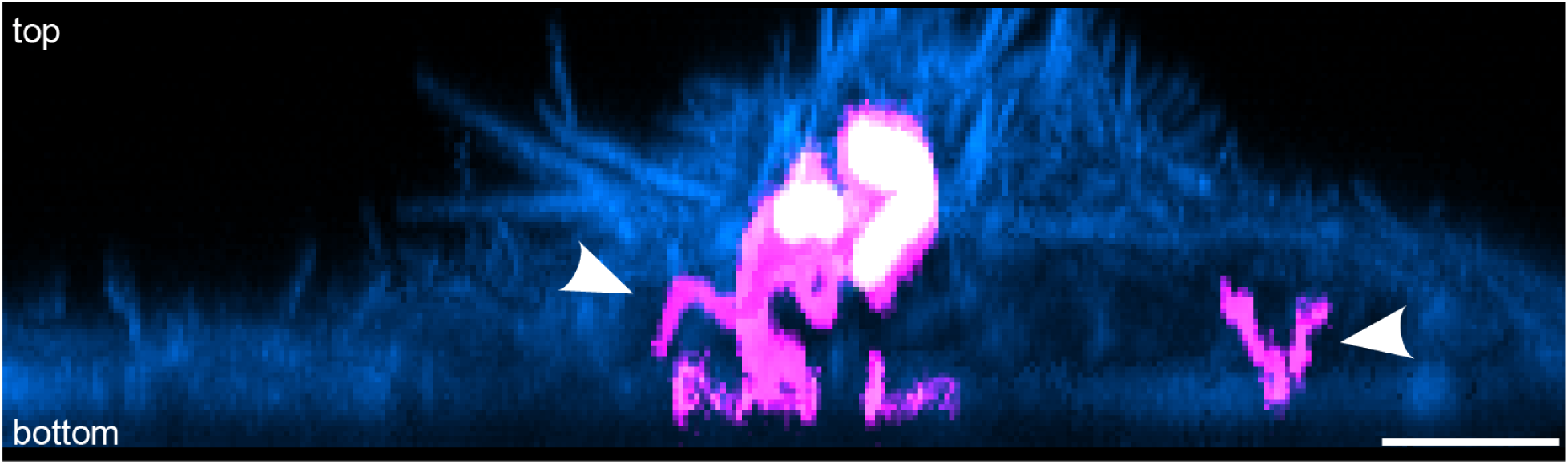
Cytosolic presence of flagella in SPA infected cells. HeLa LifeAct-GFP cells were infected with SPA FliC^E234C^, maleimide-labeled flagella and constitutive expression of mScarlet-I at MOI 30. Cells were fixed 1 h p.i. and imaged using 3D-STED microscopy. HeLa LifeAct-GFP cells are shown in cyan, flagella in magenta and SPA in white. Single flagella are marked by arrows. Scale bar, 2 µm.

**SI Appendix, Fig. S 4.**
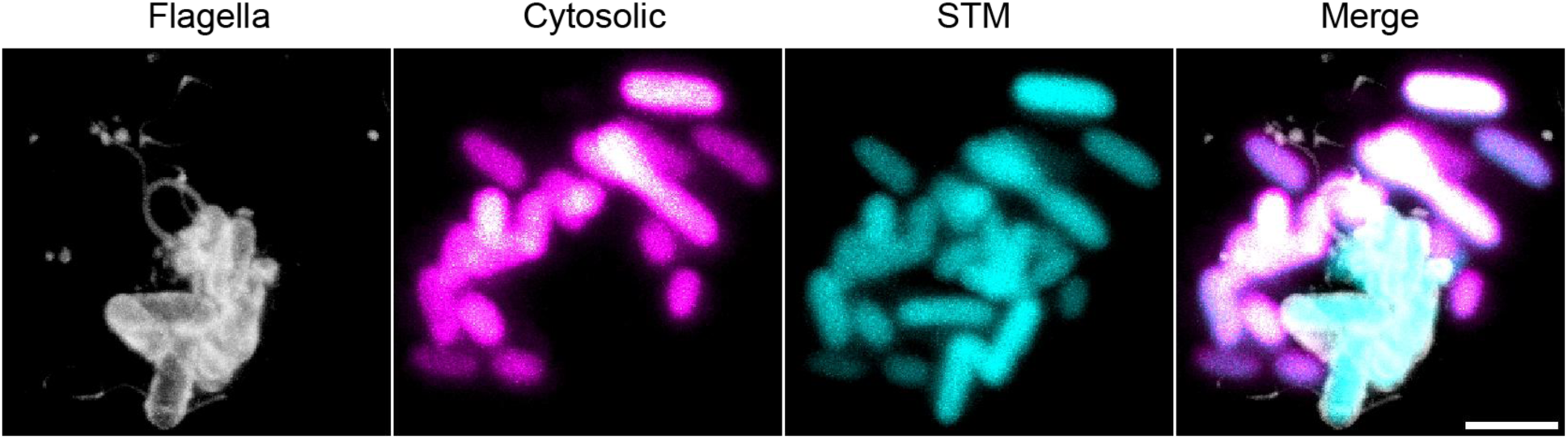
Cytosolic STM are not flagellated. HeLa WT cells were infected with STM FliC^E247C^ harboring a dual reporter plasmid with constitutive mScarlet-I (P_rpsM_-mScarlet-I) expression, a cytosol reporter (P_uhpT_-mNeonGreen-ASV) and prior to invasion labeled flagella at MOI of 30. Samples were fixed 4 h p.i. and imaged using STED microscopy. Scale bar, 2 µm.

**SI Appendix, Fig. S 5:**
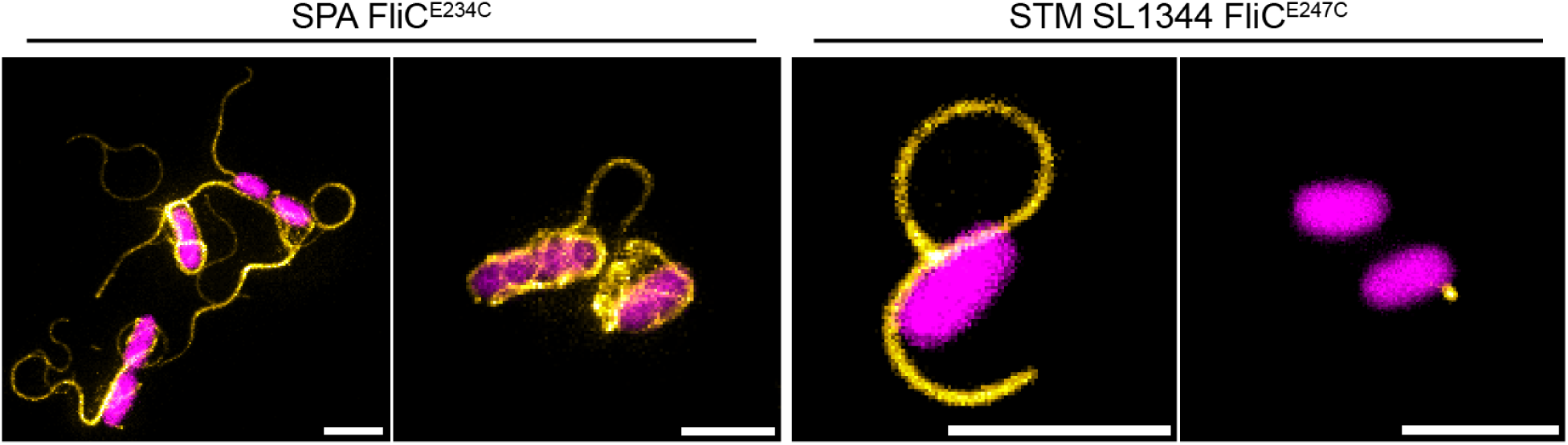
Representative comparison of intracellular flagellation of SPA and STM. STED microscopy of intracellular STM FliC^E247C^ and SPA FliC^E247C^ and prior to invasion labeled flagella and constitutive expression of mScarlet-I from plasmid. HeLa WT cells were infected with the respective serovar at MOI 30 and fixed immediately after invasion for 30 min. *Salmonella* are shown in magenta and flagella in yellow. Scale bars, 2 µm.

**SI Appendix, Fig. S 6:**
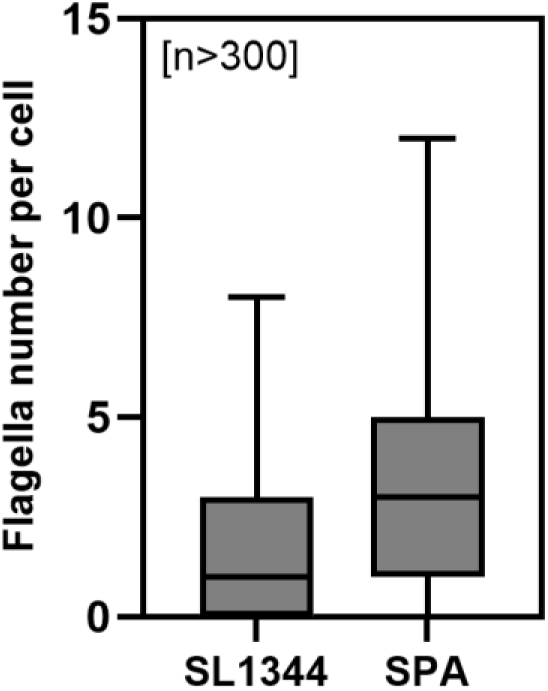
SPA shows higher flagellation than STM. Flagellation of SPA FliC^E234C^ and STM FliC^E247C^ was characterized by fluorescence microscopy after maleimide labeling of flagellar filaments. STM were grown for 3.5 h under aerobic conditions and SPA for 16 h under microaerobic conditions, respectively, before maleimide labeling. Labeled bacteria were loaded into flow cells, fixed with 4% formaldehyde, and imaged by fluorescence microscopy. Flagella numbers were quantified using ImageJ. More than 300 cells were analyzed for each strain. Data are shown as box-and-whisker plots, with boxes indicating the interquartile range, center lines indicating the median, and whiskers indicating the minimum and maximum values.

**SI Appendix, Fig. S 7:**
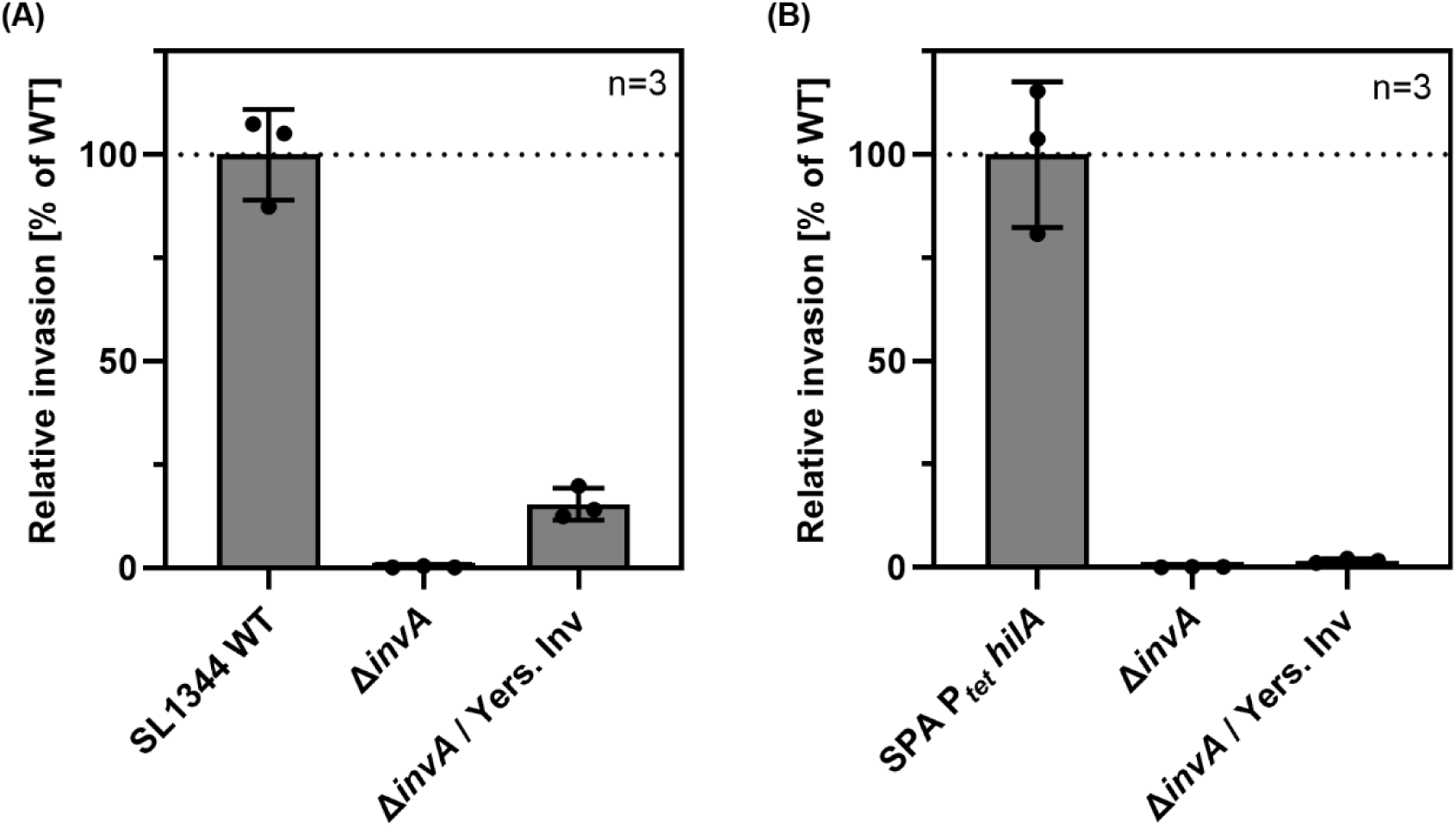
Zipper invasion complements a SPI-1 Δ*invA* mutant in STM but not SPA. Relative invasion of *Salmonella* mutants was assessed in HeLa WT cells. HeLa cells were seeded in 24-well plates to reach 100% confluence on the day of infection. Bacterial strains were subcultured from o/n cultures and grown for 3.5 h before infection. SPA (P*_tetA_*-*hilA*) was grown in the presence of 10 ng/ml AnTc. Bacteria were added to HeLa cells at an MOI of 50. Following a gentamicin protection assay with 100 µg/ml gentamicin, cells were lysed for 15 min in PBS containing 0.1% Triton X-100. Recovered intracellular bacteria and bacterial inocula were plated on LB agar plates and incubated overnight at 37 °C. Colony-forming units were quantified, and invasion was calculated relative to SL1344 or SPA (P*_tetA_*-*hilA*), respectively.

**SI Appendix, Fig. S 8:**
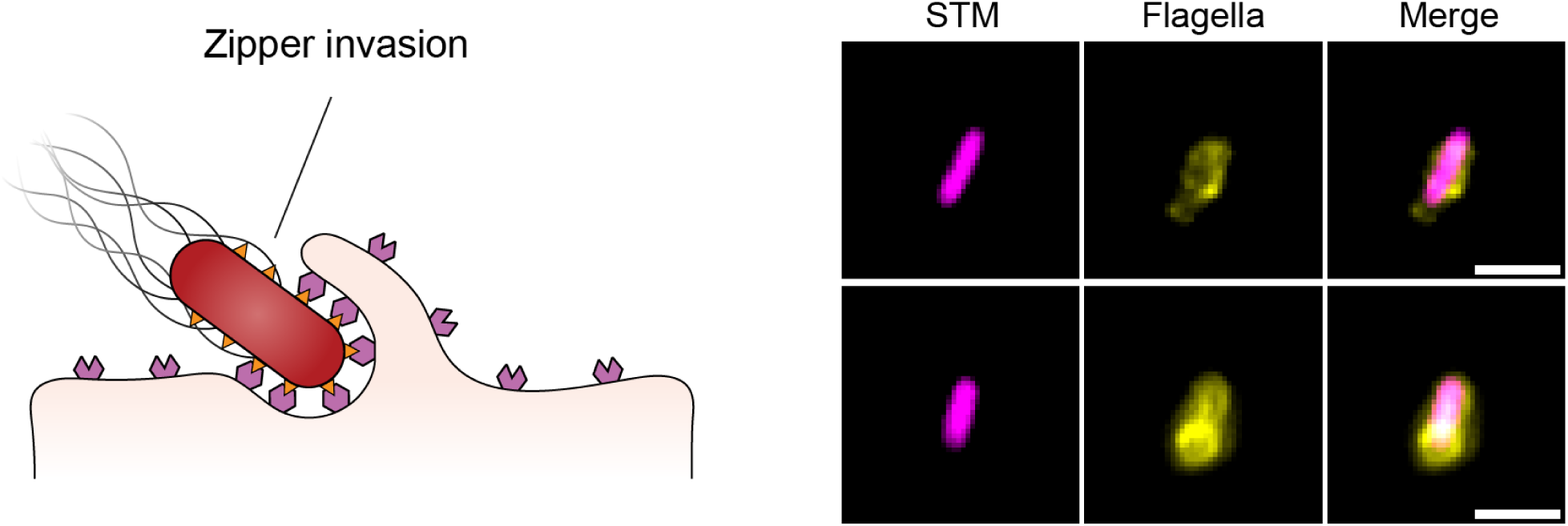
STM that invaded via zipper mechanism remain flagellated. Zipper invasion mechanism is illustrated left. Intracellular flagellation of zipper invaded STM is shown on the right. HeLa WT cells were infected with SPI-1 mutant STM FliC^E247C^ (Δ*invA*) with constitutive expression of the *Yersinia* invasin gene *inv* at MOI 100. Scale bars, 2 µm.

**SI Appendix, Fig. S 9:**
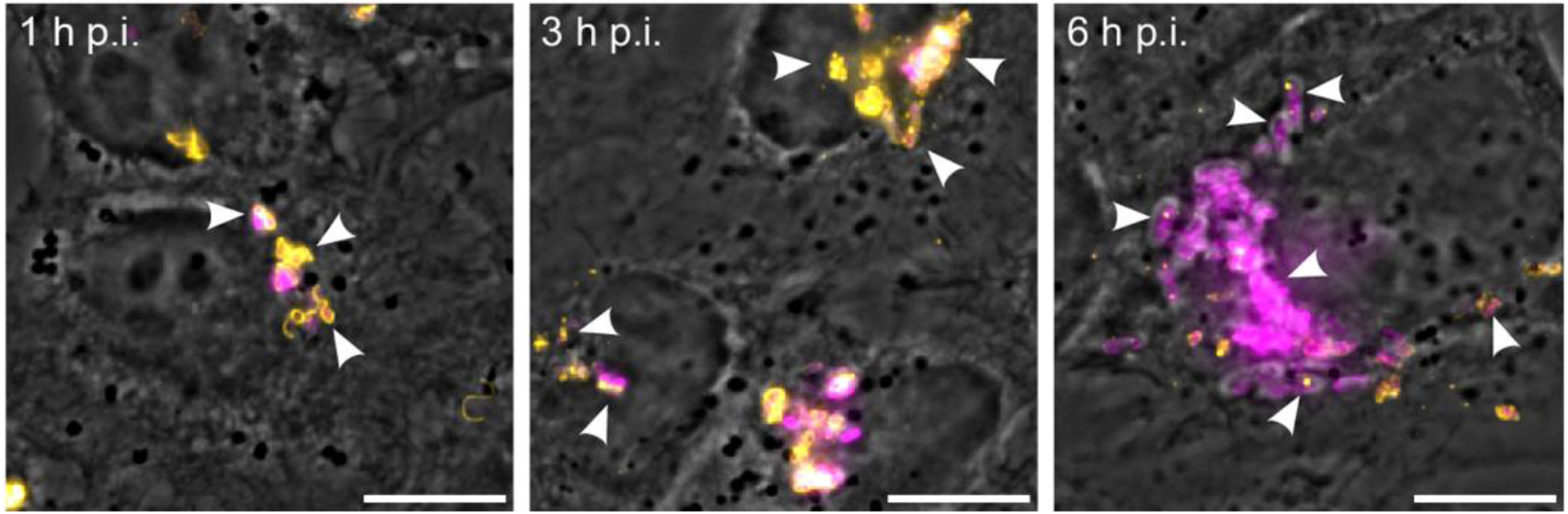
Internalized flagella disassemble over time. HeLa WT cells were infected with STM SL1344 constitutively expressing mScarlet-I from plasmid at MOI 30. After incubation for 1, 3, or 6 h p.i. samples were fixed and FljB and FliC flagellins were stained via immunostaining. Samples were imaged by phase contrast, and epifluorescence for flagella (yellow) and STM (magenta). Progressive flagella disassembly over time of intracellular presence is indicated by white arrows. Scale bars, 10 µm.

**SI Appendix, Fig. S 10:**
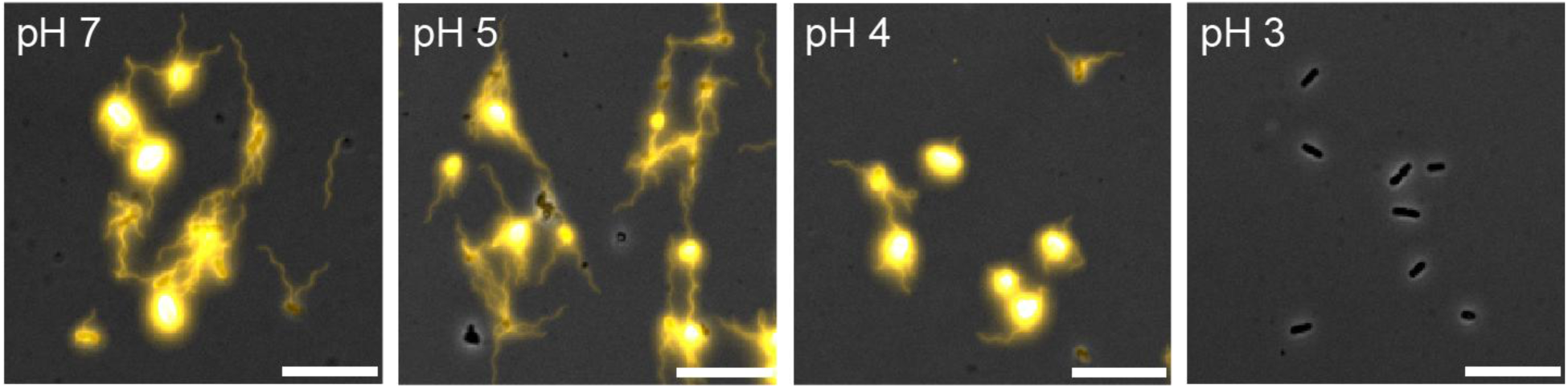
Flagella are susceptible to acidic pH. STM SL1344 fliC-ON was grown overnight, diluted 1:100 into fresh LB medium, and incubated for a further 3 h. Bacterial cultures were then transferred to buffers of different pH at an OD of 0.2 in a final volume of 1 ml. Suspensions were incubated for 2 h at 37 °C with shaking at 400 rpm in a thermo-shaker. Bacteria were subsequently centrifuged for 10 min at 2,500 × g and resuspended in PBS, pH 7. Bacterial suspensions were added to poly-L-lysine-coated flow cells, incubated for 2 min, and fixed for 5 min. FliC flagellin was immuno-stained and imaged by fluorescence microscopy. Scale bars, 10 µm.

**SI Appendix, Fig. S 11:**
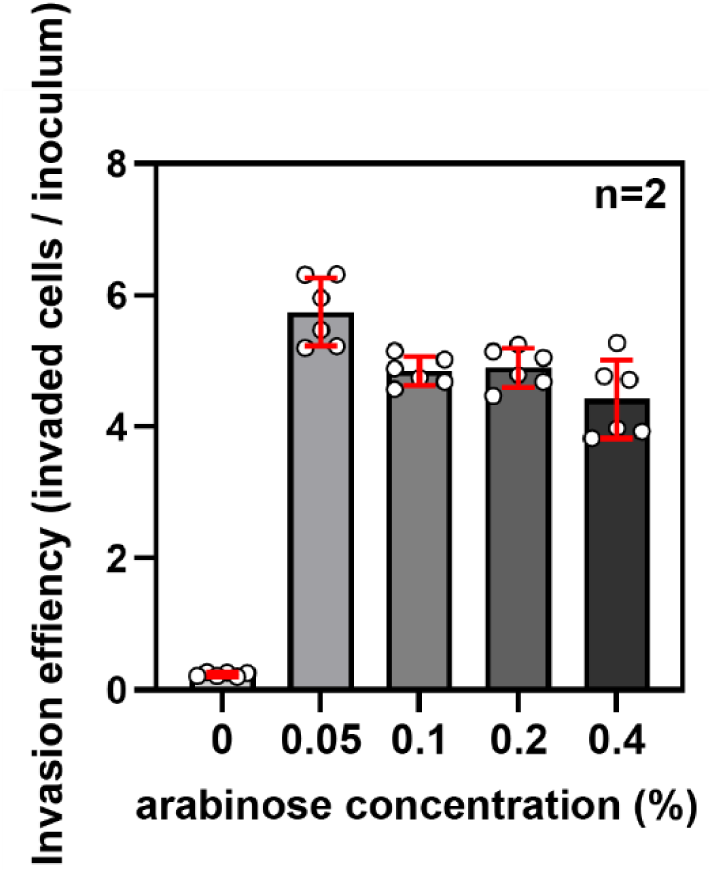
Synthetic expression of *hilA* increases invasiveness of SPA. Invasion of HeLa WT cells by SPA was assessed by infection at an MOI of 10 using SPA P*_ara_*-*hilA* constitutively expressing mScarlet-I from a plasmid. Bacterial cultures were subcultured from o/n cultures and grown for 3.5 h in LB-Miller medium containing ampicillin and supplemented with the indicated concentrations of arabinose. After killing extracellular bacteria with gentamicin, HeLa cells were detached from the wells using Accutase and fixed with PFA for 10 min. Infected HeLa cells were quantified by flow cytometry based on the mScarlet-I signal. For each replicate, 30,000 events were analyzed. The bacterial inoculum was serially diluted, plated in technical triplicates on LB agar plates containing ampicillin, and quantified by CFU counting. Invasion efficiency was calculated as the ratio of infected HeLa cells to the plated bacterial inoculum. Invasion efficiency was determined from two biological replicates, each consisting of three technical replicates, and is shown as the mean ± SD in red.

**SI Appendix, Fig. S 12:**
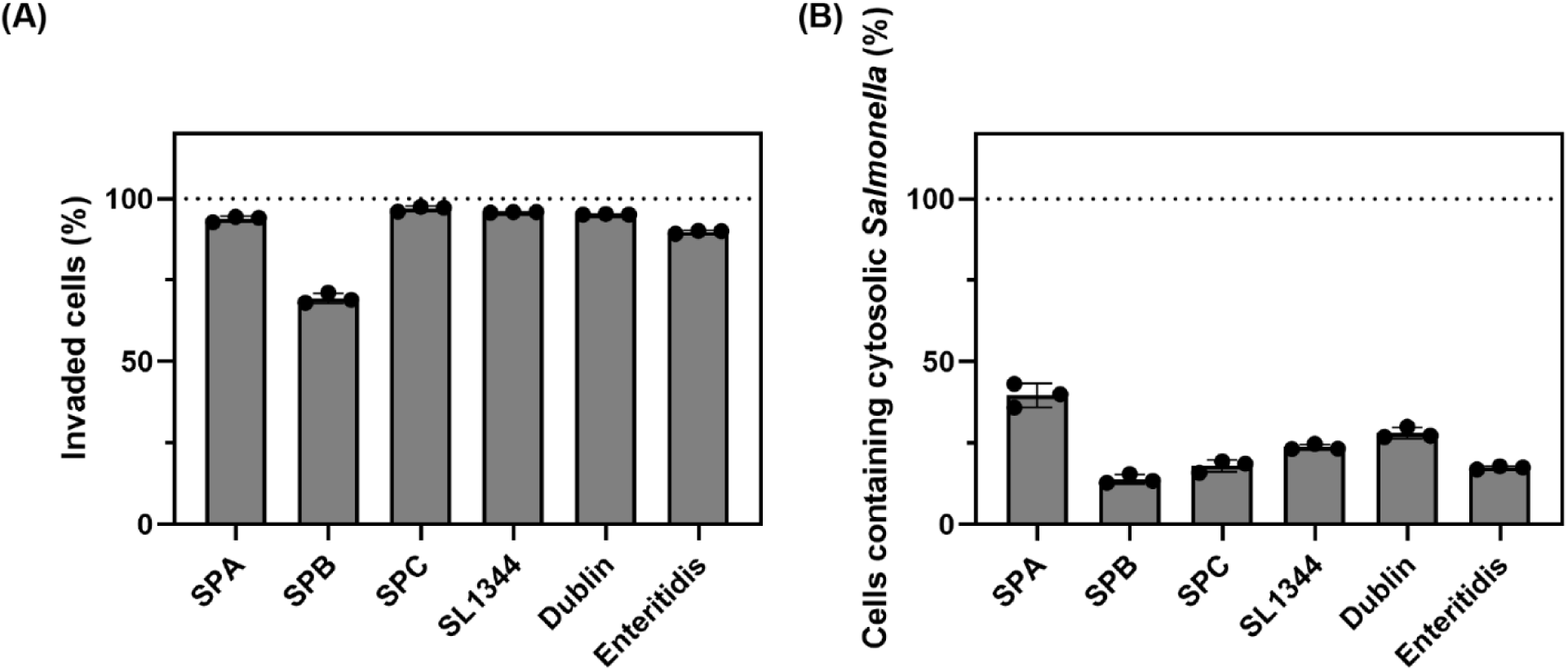
Cytosolic subfraction of *Salmonella* isolates. The cytosolic subfraction of *Salmonella* isolates was determined in HeLa WT cells. Bacteria were transformed with a dual-reporter plasmid enabling detection of both cytosolic bacteria and total intracellular bacteria through a constitutive signal. Bacterial cultures were subcultured from o/n cultures and grown for 3.5 h in LB-Miller medium containing chloramphenicol and supplemented with 0.05% arabinose for SPA P*_ara_*::*hilA*. After killing extracellular bacteria with gentamicin, HeLa cells were detached from the wells using Accutase and fixed with 4% formaldehyde for 10 min. Invasion **(A)** and cytosolic subfraction **(B)** were quantified by flow cytometry and are shown as mean values from three biological replicates with SD. For each replicate and strain, 30,000 events were analyzed to quantify both invaded cells and cells containing cytosolic bacteria.

**SI Appendix, Fig. S 13:**
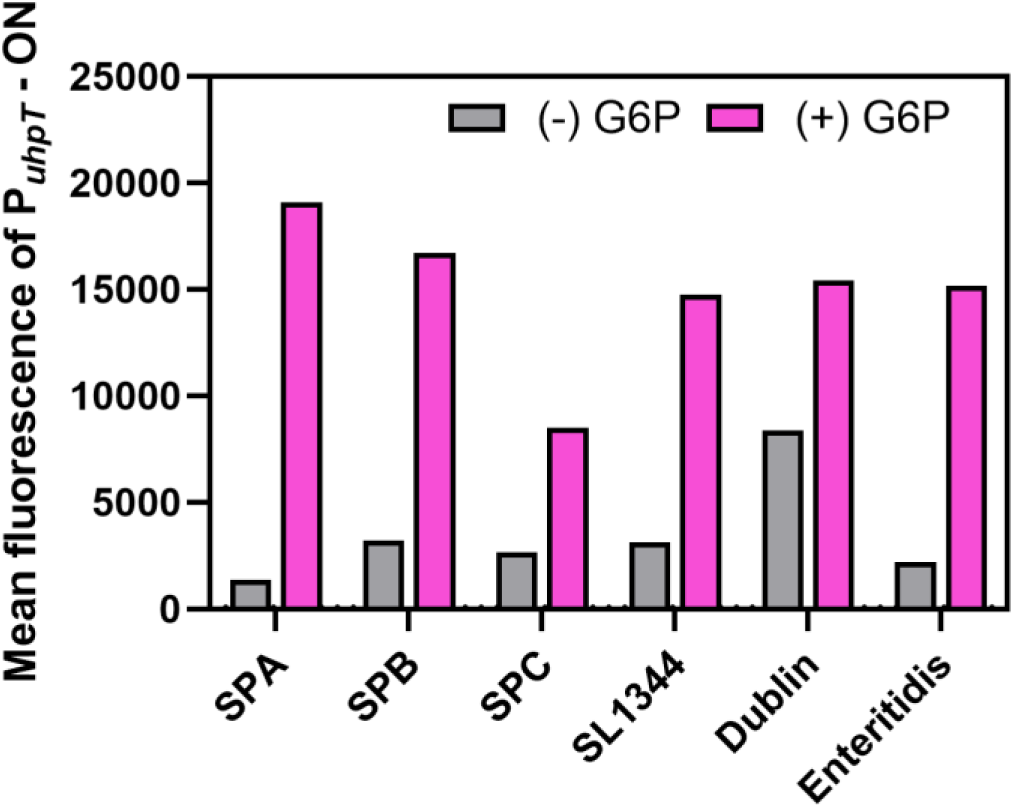
Performance of the cytosolic reporter in various *Salmonella* serovars. Reporter signal strength was assessed in different *Salmonella* serovars. Strains carrying the dual-reporter plasmid were grown o/n in LB supplemented with ampicillin, diluted 1:100 into 3 ml fresh LB in the presence or absence of 0.2% G6P, and grown for 2 h at 37 °C with shaking. The OD of each culture was adjusted to 0.1, and samples were fixed with 4% formaldehyde for 5 min. Cells were washed twice in PBS and analyzed by flow cytometry. The mScarlet-I signal was used to distinguish bacteria from debris, and the mNeonGreen signal was used to assess cytosolic reporter responsiveness in each serovar. The mean mNeonGreen fluorescence of the population is shown. More than 50,000 events were analyzed for each condition (n > 50,000).

**SI Appendix, Table S 1:**
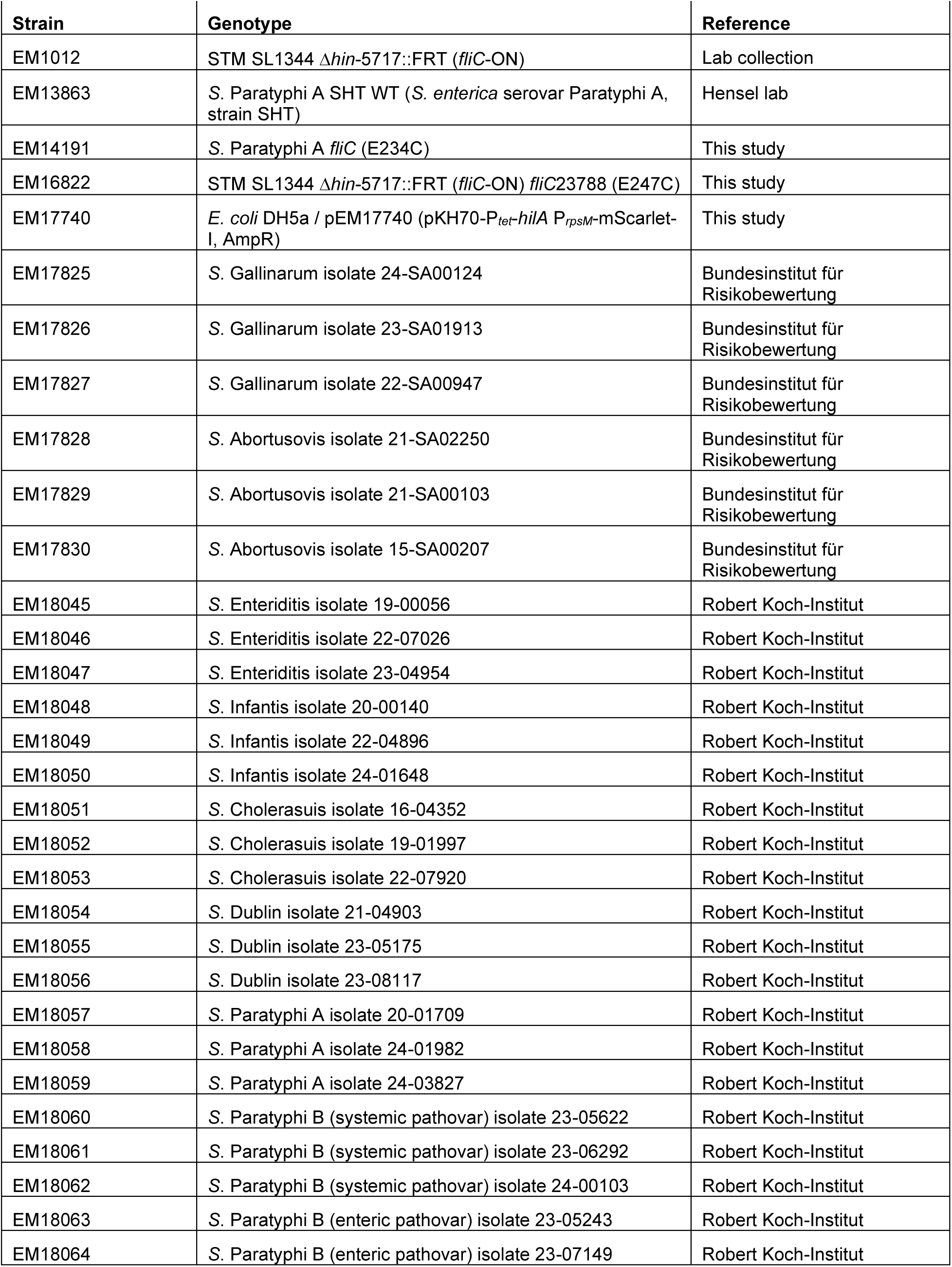
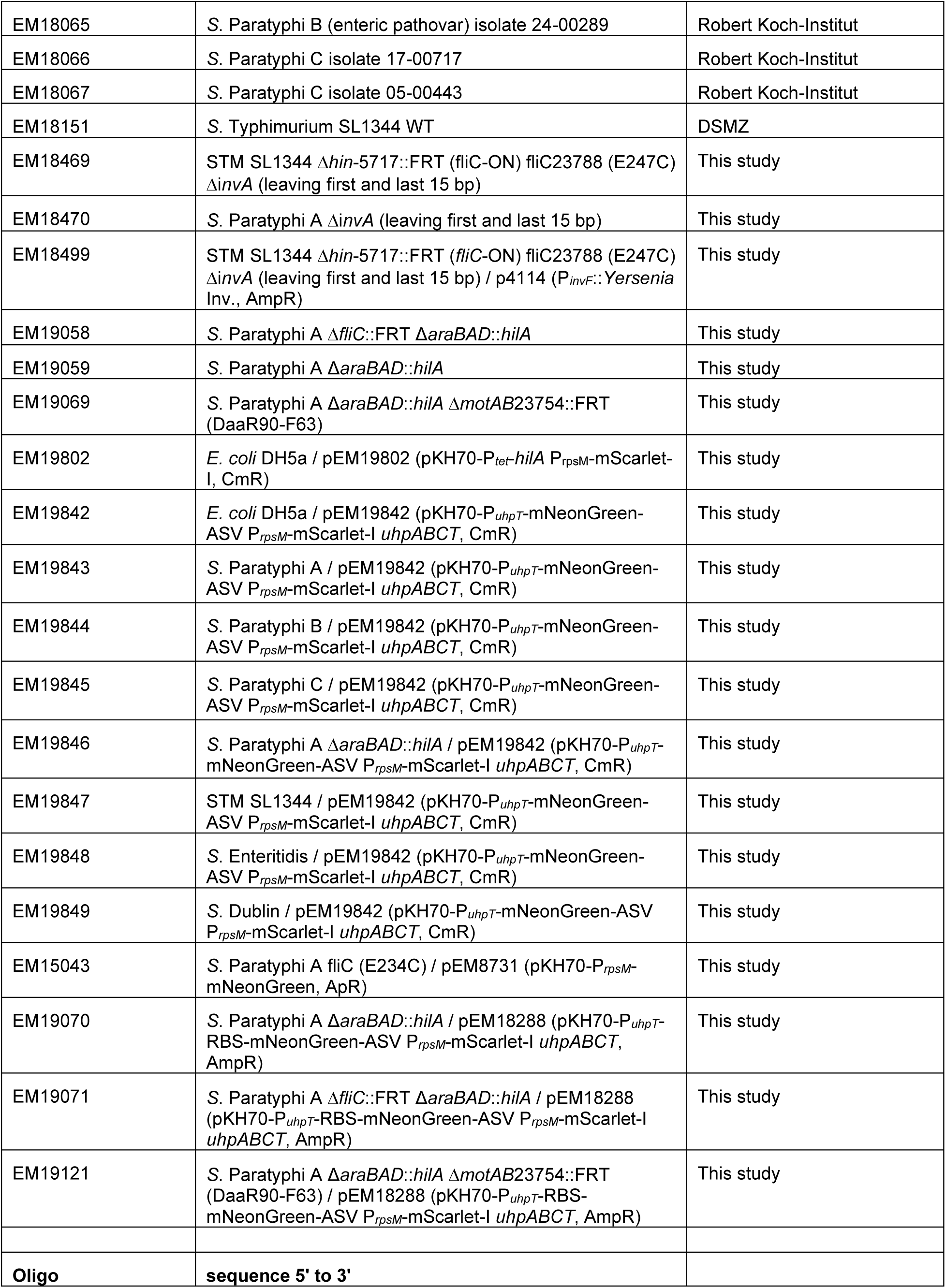
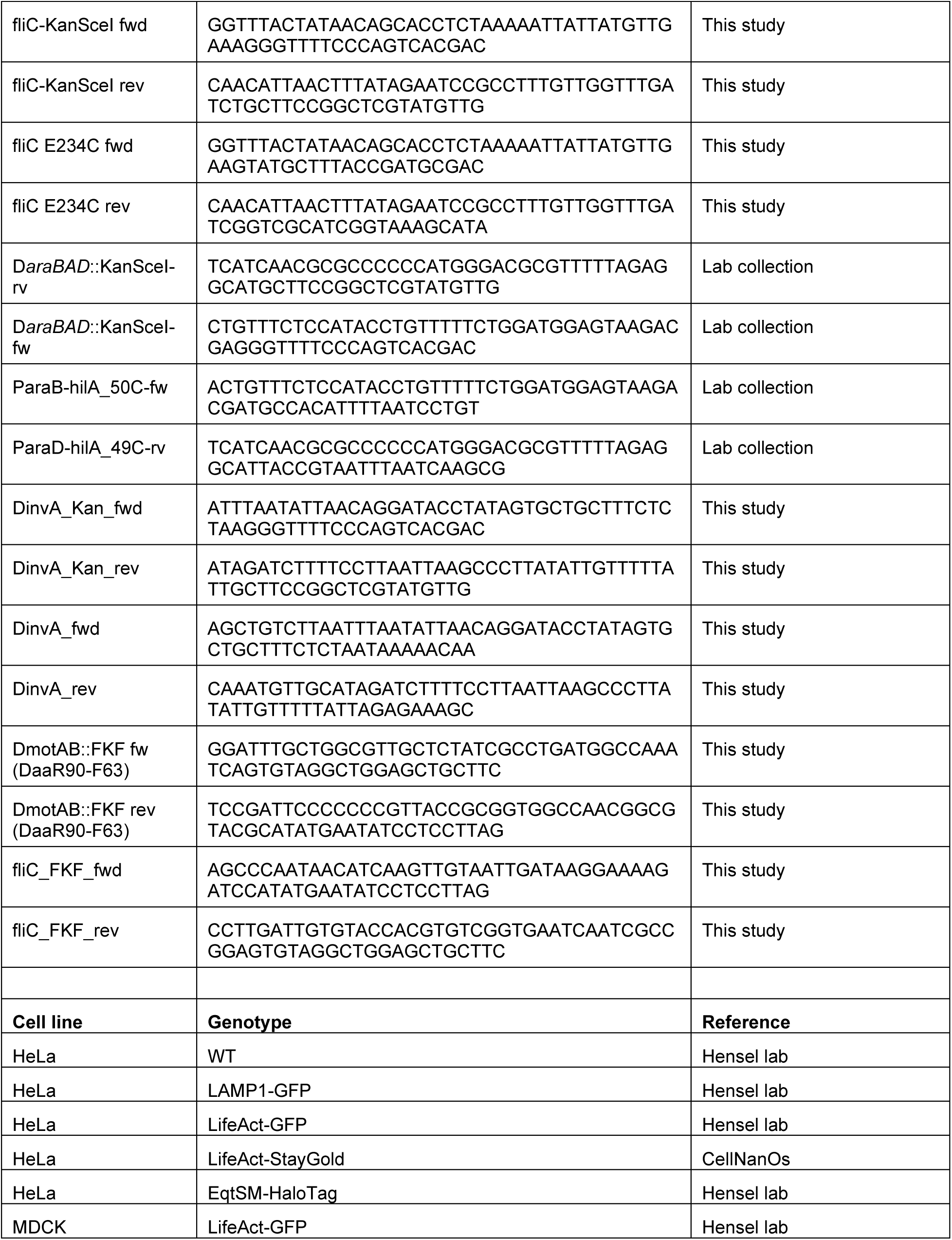
Strains, oligonucleotides used in this study.

## Supporting movie information

**Movie S1, S2 and S3: LLSM of STM FliC^E247C^ invading into HeLa LifeAct-StayGold cells.** Shown are STM constitutively expressing mScarlet-I (magenta), maleimide-labeled flagella (yellow), and HeLa LifeAct-StayGold cells (cyan). Bacteria were diluted 1:31 from an o/n culture into LB-Miller supplemented with ampicillin and grown for 3.5 h in fresh LB. The OD_600_ of bacterial cultures was adjusted to 0.1 in 1 ml PBS, and flagella were stained with 10 µM maleimide StarRed for 5 min. After one wash in PBS, 100 µl of the bacterial suspension was directly added to the sample bath. Time-lapse videos were further processed in Imaris.

**Movie S4: Representative LCI for intra-SCV motility phenotype of SPA:** Shown are SPA constitutively expressing mScarlet-I (magenta), maleimide-labeled flagella (yellow), and HeLa LAMP1 cells (cyan) at 2 h p.i. Bacteria were cultured under microaerobe conditions for 16 h in LB-Miller supplemented with ampicillin. The OD_600_ of bacterial cultures was adjusted to 0.2 in 1 ml PBS, and flagella were stained with 10 µM maleimide StarRed for 10 min. HeLa cells were infected at MOI 30. Scale bar, 10 µm.

**Movie S5: LLSM time-lapse movie of HeLa EqtSM cells infected with SPA.** HeLa EqtSM cells are shown in cyan, SPA constitutively expressing mScarlet-I in magenta, and maleimide-labeled flagella in yellow. Bacteria were cultured under microaerobic conditions for 16 h in LB-Miller supplemented with ampicillin. The OD_600_ of bacterial cultures was adjusted to 0.2 in 1 ml PBS, and flagella were stained with 10 µM maleimide StarRed for 10 min before addition to the cells. To enhance contact with host cells, bacteria were centrifuged onto HeLa cells for 5 min at 500 × g.

**Movies S6 and S7. Representative movies of intracellular flagella-mediated motility.** HeLa LAMP1 cells are shown in cyan, SPA in magenta, and labeled flagella in yellow at 2–3 h p.i. Bacteria were cultured under microaerobic conditions for 16 h in LB-Miller supplemented with ampicillin. The OD_600_ of bacterial cultures was adjusted to 0.2 in 1 ml PBS, and flagella were stained with 10 µM maleimide StarRed for 10 min. HeLa cells were infected at MOI 30. Scale bar, 10 µm.

